# Striatal indirect pathway mediates action switching via modulation of collicular dynamics

**DOI:** 10.1101/2020.10.01.319574

**Authors:** Jaeeon Lee, Bernardo L. Sabatini

## Abstract

Type 2 dopamine receptor-expressing, or indirect pathway striatal projection (iSPNs), neurons comprise one of two major pathways through the basal ganglia^1^, and are a major drug target for treatment of neuropsychiatric disorders^2–4^. The function of iSPNs is unclear with proposed roles in suppression of unwanted actions and in refining selection actions or their kinematics^5–12^. Here, we show that iSPNs can simultaneously suppress and facilitate conflicting motor actions in a lateralized licking task. Activation of iSPNs suppresses contraversive while promoting ipsiversive licking, allowing mice to rapidly switch between alternative motor programs. iSPN activity is prokinetic even when mice are not cued to perform an action. Activity in lateral superior colliculus (lSC), a basal ganglia target, is necessary for performing the task and predicts action. Furthermore, iSPN activation suppresses ipsilateral lSC, but surprisingly, excites contralateral lSC. iSPN activity has neuron-specific effects that, at the population level, steers the neural trajectory towards that associated with ipsiversive licking. Thus, our results reveal a previously unknown specificity of iSPNs effects on downstream brain regions, including the ability to excite contralateral regions and trigger motor programs. These results suggest a general circuit mechanism for flexible action switching during competitive selection of lateralized actions.

## Introduction

The basal ganglia (BG) is a phylogenetically old and evolutionary conserved set of brain structures implicated in action selection and motor control^13–15^. An influential model of BG function posits that Type 1 dopamine receptor (D1R)-expressing direct pathway striatal projection neurons (dSPNs) promote, whereas the Type 2 receptor (D2R)-expressing indirect pathway striatal projection neurons (iSPN) suppress movement^1,5,16,17^. Although previous studies have generally supported this view with respect to the direct pathway, controversies exist regarding the function of the indirect pathway. Alternative models posit that iSPN activity promotes the switching of actions, suppresses unwanted movements, or refines the choice of selected action^10,11,18–23^. Here, we measured behavioral and neural responses after transiently activating the indirect pathway of the BG licking circuit in mice preforming a lateralized licking task. We show that the behavioral effects of iSPN stimulation can be broken down into two seemingly independent components of contraversive movement suppression and ipsiversive movement initiation. Further, indirect pathway stimulation caused a push-pull modulation of the lateral superior colliculus (lSC), a region downstream of BG that is necessary for and is preferentially active during contraversive licking. iSPN activation suppresses and excites the ipsilateral and contralateral lSC, respectively, in a manner that depends on the action-direction preference of each neuron. At the network level, iSPN stimulation moves lSC population activity along the choice-encoding dimension in the ipsiversive direction, but has only minimal effects along other behavior-related dimensions ^24^. Our results suggest a general circuit model in which simultaneous suppression and activation of ipsilateral and contralateral lSC via the indirect pathway facilitates action switching and thus permits exploration of alternative options during lateralized action selection.

## Results

### iSPN activity biases action selection

To manipulate iSPN activity, we expressed a Cre-dependent activator opsin (CoChR)^25^ into striatum and implanted two tapered optical fibers^26–28^ in *Adora2a-Cre* mice^29^ (**Fig. 1a, Extended Data Fig. 1a**). Tapered fibers allowed focal activation of distinct striatal regions allowing us to functionally map lick-related regions of striatum^30^. Mice were trained on a lateralized licking task during which a brief (50 ms) auditory cue (tone A or B) indicated the spout they had to lick in order to receive a water reward (tone A->lick left; tone B->lick right, **Fig. 1b**). We categorized the outcome of each trial, based on the timing and direction of the first lick, as either correct, incorrect, or miss (no licks within 500 ms after tone onset). If the trial outcome was correct, a solenoid opened to deliver a water reward. We refer to the first lick is referred to as the “choice” lick. After two weeks of training, all mice achieved above 79% correct rate (mean correct rate = left: 93%; right: 95%), with a latency of on average ~130 ms from the tone onset to the choice lick (**Extended Data Fig. 1b, c**).

**Fig. 1.**
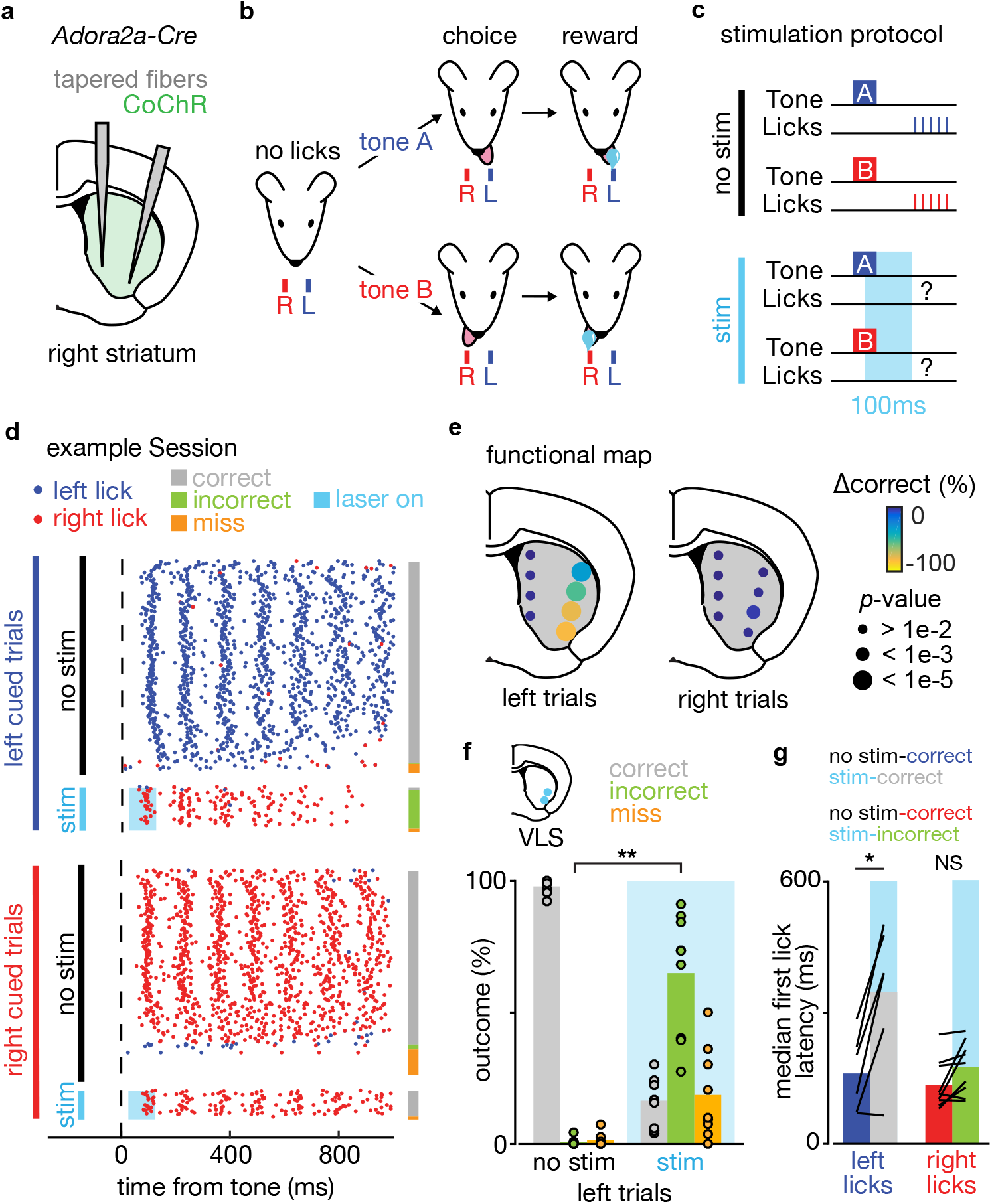
Indirect pathway activation in a striatal subzone biases towards ipsiversive licking. **a,** Schematic coronal section showing CoChR expression in the striatum (green) of an *Adora2a-Cre* mouse with two tapered fibers (gray) implanted in medial and lateral striatum. **b.** Task structure: a head-fixed mouse withholds licking (no lick) for a period after which a tone is played whose frequency indicates the rewarded port (left or right) upon licking correctly on the first ‘choice’ lick. **c.** Stimulation protocol: the laser (light blue) is turned on in a subset of trials after tone onset (25 ms delay) for 100 ms to test the effect on lick decision. **d.** Results from a representative session: each row shows behavior in a single trial with each dot representing a lick to the left (blue) or right (red) port relative to tone onset. Trials are sorted by trial type (top: left cued trials, bottom: right cued trials) and further divided into no stimulation trials (black) and optogenetic stimulation trials (light blue). The far-right column shows the trial outcomes labeled as correct (gray), incorrect (green), or miss (orange). **e.** Map of functional perturbations summarized as the change in percentage of correct trials (Δcorrect) induced by optogenetic stimulation on left- or right-cued trials. Each circle indicates a striatal stimulation site (total 8 sites). The color and size of each circle denote the effect size and p-value (bootstrap, see Methods), respectively (n=5 mice, 9 sessions). Stimulation in the ventrolateral striatum (VLS) was the most effective site at perturbing performance (Δ_correct_=-82%, P<1e-5). **f.**Quantification of trial outcomes resulting from VLS stimulation (n=9 sessions): percentages of correct (grey), incorrect (green) and miss (orange) outcomes in no stimulation (left) and stimulation trials (right). Stimulation caused a significant increase in percentage of incorrect trials (P**<0.001). **g.** Median first lick latency: licks in no stimulation trials are separated into left vs. right (blue/red) whereas those during stimulation trials are sorted into incorrect vs. correct (green/grey). Correct licks during stimulation trials to the left were delayed compared to correct licks during no stimulation trials (P*<0.05, two-tailed *t*-test) (left licks: n= 6 sessions, right licks: n=9 sessions, see Methods).

To determine which regions of the striatum iSPN activity was able to perturb task performance, we stimulated iSPN in distinct striatal regions (8 sites, 4 sites per fiber) on the right side of the brain. Optogenetic stimuli were delivered after the start of the instructive cue known as the ‘decision window’ (**Fig. 1c**). Brief unilateral iSPN stimulation immediately after tone onset (100 ms pulse starting 25 ms after tone onset delivered in randomly interspersed ~30% of trials) in trained mice during this decision window decreased the fraction of correct outcomes (**Fig. 1d, e**). This effect was only present in left-cued trials (contralateral to the fiber location) and was specific to stimulation of the ventrolateral striatum (VLS) (**Fig. 1e, Extended Data Fig. 1d**). VLS stimulation induced error trials consisting largely of incorrect choice licks as opposed to misses (**Fig. 1f**). The outcomes of left-cued trials following the stimulation trial were unaffected, indicating that the stimulation protocol did not cause a persistent change in behavior or action value (**Extended Data Fig. 1e**). When mice selected the correct spout despite the stimulation, licks were delayed relative to those in control trials, suggesting an additional interaction between the stimulation and the motor program to lick to the correct spout (**Fig. 1g**).

### iSPN activation induces a motor action

Given the hypothesized function of iSPN in movement suppression^5,16^, it is surprising that following iSPN stimulation, results in incorrect performance on ipsiversive licks as opposed to suppression of licking all together. One possibility is that the stimulation suppressed the drive to lick to contra side, but not the general drive to lick, thus causing mice to resort to the alternative option of licking to the ipsilateral spout. Another possibility is that iSPN stimulation itself promoted licking to the ipsilateral side. This model makes the prediction that iSPN stimulation will induce ipsiversive licking independent of planning a movement to the contralateral side.

In order to test these two models, we designed an extinction paradigm in which we initially trained mice on the same lateralized licking task (**Fig. 1b**), but subsequently devalued the right spout by not delivering reward to it even after correct spout selection (**Fig. 2**). Stimulating iSPN before and after extinction allowed us to compare the effect of removing the drive to lick the right spout and thus determine if the iSPN stimulation can generate ipsiversive licking only when a contraversive movement is being suppressed. We bilaterally implanted tapered fibers, one targeting VLS in each striatal hemisphere, in *Adora2a-Cre* mice expressing CoChR. The mice were trained to perform the main task, and optogenetically stimulated as before, but alsoafter extinction of the right spout (**Fig. 2a, c**). Given that stimulation generally only affected contraversive trials, we focused our analysis on trials in which the cue indicated that the correct choice lick was to the spout contralateral of the stimulation site (**Fig. 1e**).

**Fig. 2.**
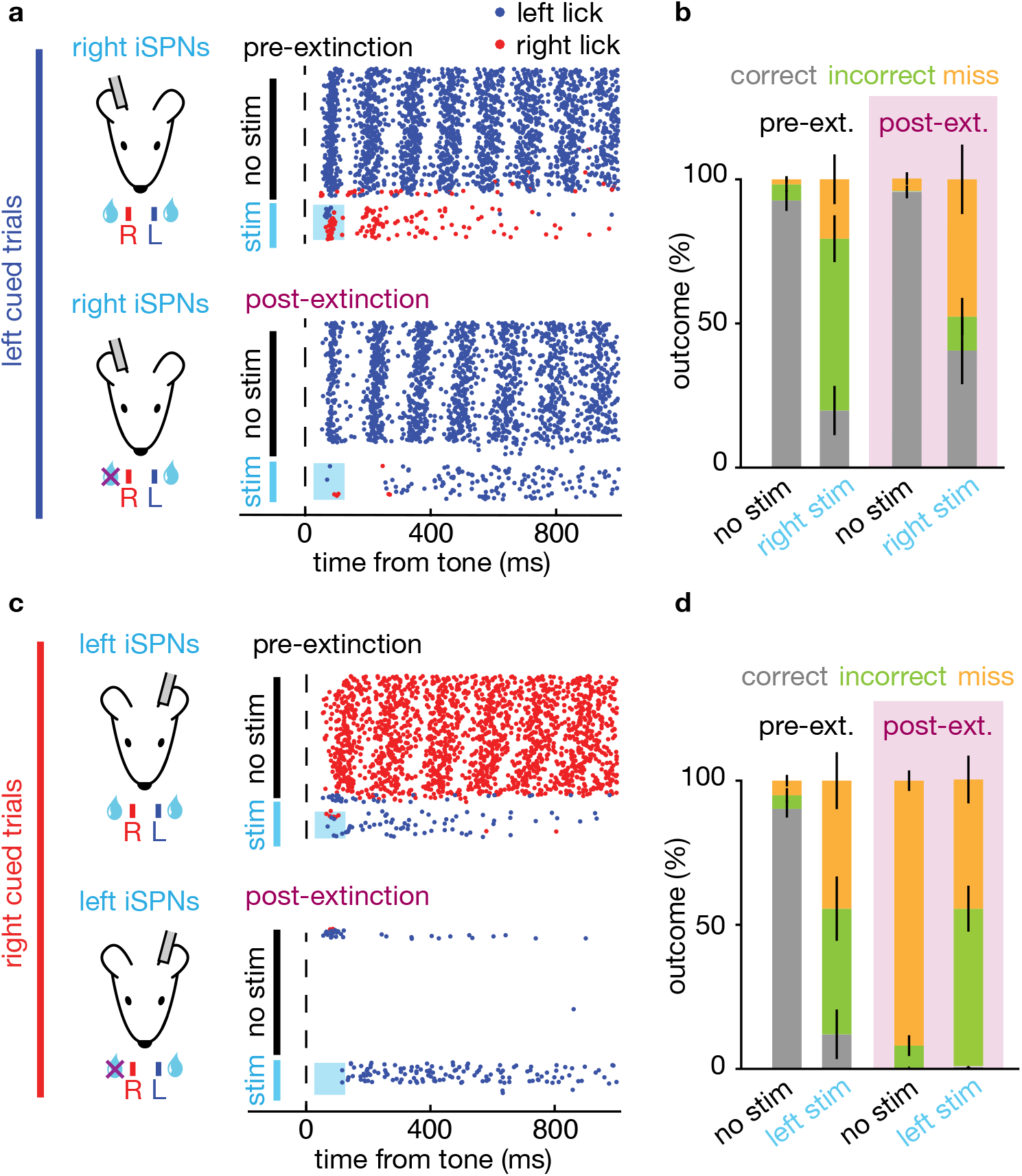
iSPN activation induces ipsiversive licking independent of contraversive lick suppression. **a, c.** An example session from one mouse showing the effects of iSPNs stimulation on the right (**a**) or left (**c**) hemisphere before (pre-extinction, top) and after (post-extinction, bottom) devaluation of the right port. Each dot represent licking either to the left (blue) or right (red) and trials (rows) are sorted by no stimulation (black) and stimulation trials (light blue). Only trials with licking cued to the port contralateral to optogenetic stimulation (left in **a**, right in **c**) are shown. **b, d.** Percentages of each outcome types for pre-extinction (black, left) and post-extinction (purple, right) optogenetic stimulation trials (stim, light blue) and control trials (no stim, black). Outcomes are color-coded grey (correct), green (incorrect), and orange (miss) (n=5 mice). The selection of the incorrect port following optogenetic stimulation of iSPNs on the right striatum significantly decreased after extinction (P<0.0125, one-tailed *t*-test), whereas it remained the same for iSPNs stimulation on the left (P=0.65, one-tailed *t*-test).

We first compared the effects of right iSPN stimulation before and after extinction of the right lick motor program. Pre-extinction stimulation caused an increase in incorrect licks to the right (ipsilateral) side, consistent with our previous results (Fib 1d and **Fig. 2a, b**). However, post extinction stimulation in the right VLS no longer caused mice to lick incorrectly to the devalued (right, ipsilateral) spout, and instead increased the fractions of miss and correct trials (**Fig. 2b**). This suggests that iSPN simulation causes mice to switch to an alternative ipsiversive motor program (licking the ipsilateral spout) only if the latter is a valuable option. We next analyzed the effect of stimulating iSPN in the left VLS. Pre-extinction stimulation lead to incorrect licking, as reported above (**Fig 1d** and **Fig. 2c**). After extinction in no-stimulation trials, mice no longer licked to the devalued spout (right trials), causing an increase in the fraction of miss trials (**Fig. 2c**) and consistent with the mouse having devalued this motor action. Surprisingly, iSPN stimulation after extinction still caused mice to lick to the ipsilateral spout (**Fig. 2c-d**). This suggests that ipsiversive licking triggered by iSPN stimulation is not a consequence of suppressing licking to the contralateral side and might reflect an ability of iSPN to directly trigger a learned motor action. Bilateral iSPN stimulation increased miss rate but not incorrect rate, suggesting the phenotype is a unique consequence of unilateral stimulation (**Extended Data Fig. 2a**). We also trained a separate group of mice to only lick to the left spout (see Methods). Right iSPN stimulation in these mice failed to induce licking of the right spout, suggesting that stimulation induced licking is not a hardwired motor program (**Extended Data Fig. 2b-c**). In mice trained on the main two-spout task (**Fig. 1b**), we also observed that iSPN stimulation during the inter-trial-interval when mice rarely licked, induced ipsiversive licking although this effect emerged over multiple stimulation sessions, suggesting iSPN have the capacity to generate a motor program by itself outside the decision window (**Extended Data Fig. 3; see Methods**). Overall, our results support a model whereby unilateral iSPN activation can cause ipsiversive licking, even in the absence of contraversive licking suppression, but only if it is a reinforced motor action.

### lSC activity drives contraversive licking and predicts upcoming choice

Superior colliculus (SC) is a region downstream of BG implicated in decision making and behavioral competition^6,31–36^. Activity in the lateral part of SC (lSC) is necessary for licking behavior^37^. Thus, we hypothesized that the behavioral phenotype induced by iSPN stimulation might arise from modulation of activity in lSC. We first asked if BG output innervates lSC by examining the axons of SNr neurons downstream of VLS, labelled via AAV1-Cre mediated anterograde tracing (**Fig. 3a**) (Lee *et al,* manuscript in prep). VLS recipient SNr (^VLS^SNr) cells innervated lSC on both sides of the brain (contralateral and ipsilateral SC) (**Fig. 3b-c**). Interestingly, this bilateral projection was strongest in lSC-targeting ^VLS^SNr axons but much weaker in other brain regions (**Extended Data Fig. 4**). To test the potential involvement of lSC in the licking task, we infused muscimol, a GABA_A_ receptor agonist, into lSC unilaterally while mice were performing the task (see Methods). Muscimol infusion reduced the performance only on trials in which the correct selection port was contralateral to the infusion site, suggesting lSC is necessary for contraversive licking (**Fig. 3d**).

**Figure 3.**
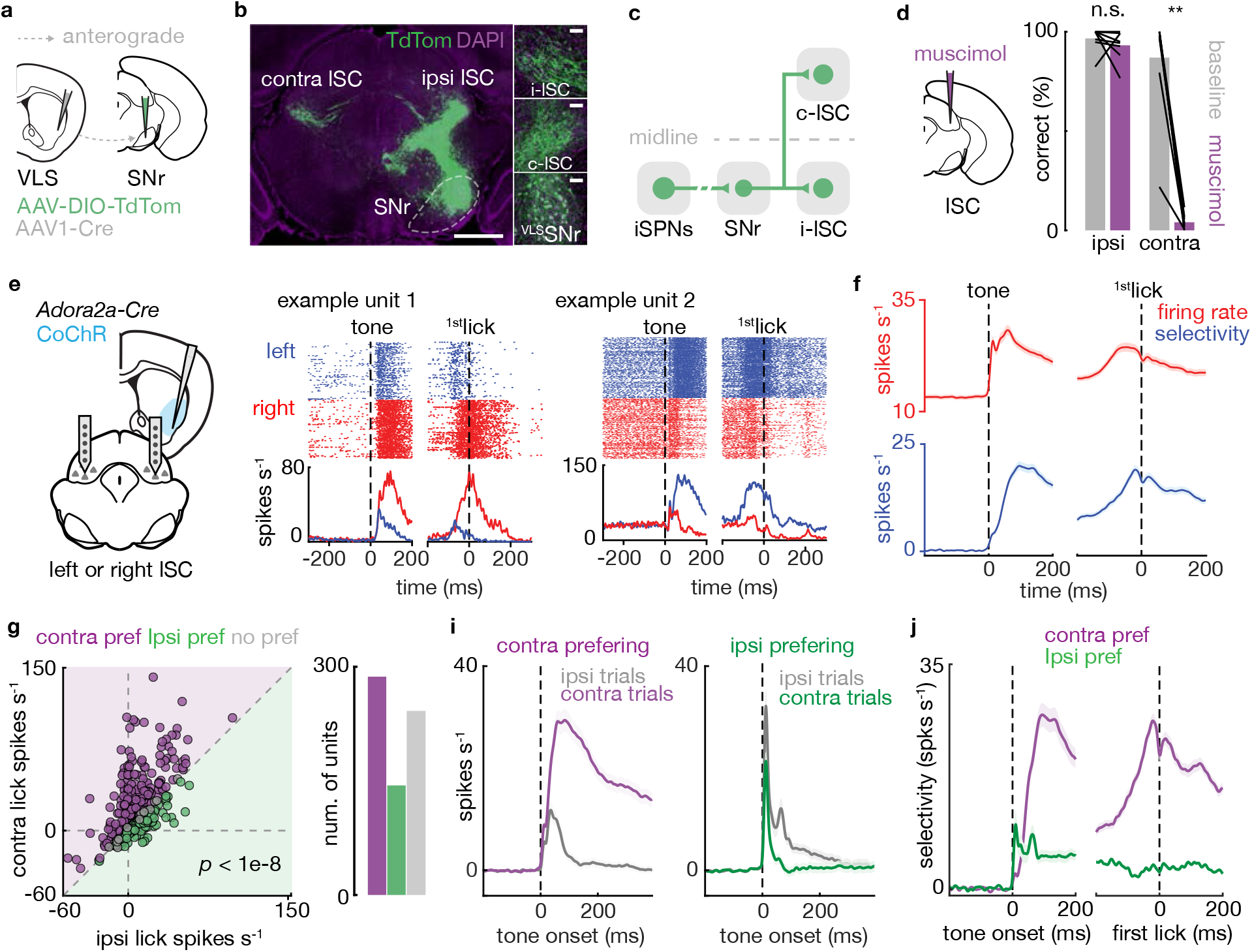
Activity in the lateral superior colliculus drives contraversive licking and predicts upcoming choice direct. **a.**Anterograde mapping: Strategy to map VLS recipient SNr (^VLS^SNr) projection using AAV1-Cre anterograde trans-synaptic movement (see Methods). **b.**Example histology of superior colliculus: ^VLS^SNr (green) projects to both ipsilateral lSC (i-lSC) and contralateral lSC (c-lSC). SNr is shown in white dotted line. Scale bar, 1mm (left panel), 100um (3 insets in right column). **c.** Circuit schematic showing the iSPN indirect projection to SNr (intermediate GPe/STN are not shown) which innervates lSC on both hemispheres. **d.** *left,* Muscimol was infused unilaterally in lSC as the mouse performed the task performance. *righ/,* percentages of correct trials before (baseline, grey) and after (muscimol, purple) infusion. Muscimol infusion significantly impaired performance of contralateral cued trials (n=8 lSC sites, 4 mice, P**<1e-6, two-tailed *t*-test). **e.** *left,* Schematic of extracellular recordings in lSC on either side of the brain (separate sessions) in mice performing the task. *right,* Peri-stimulus histogram of activity of two example units. Correct left (blue) and right (red) cued trials are shown, aligned to either tone onset or the 1^st^ (i.e. choice) lick (dashed lines). **f.** Firing rate (red) and firing rate selectivity (preferred – non-preferred, blue) aligned to tone onset and 1^st^ lick (dashed lines). Solid line shows the average and shaded areas the SEM across units (n=673 units; 7 mice) **g.** *left,* Each dot shows the average activity of one unit in the first 200 ms after tone onset (spikes/s) during contraversive trials plotted versus that in ipsiversive trials. The directional selectivity of each unit is color-coded (purple: contra; green: ipsi; grey: no preference). Overall population activity was higher during contraversive trials (P<1e-8, two-tailed *t*-test). *right,* Numbers of cells preferring contra, ipsi trials, or no preference. **h.** Average activities of contraversive- (purple) and ipsiservice- (green) preferring units shown aligned to tone onset (dashed line) during contralateral and ipsilateral cued trials (contra: n= 296, ipsi: n=139; mean ± s.e.m. across units). **i.** Selectivity (spikes/s; activity in preferred - antipreferred trials) aligned to tone onset (left) or 1^st^ lick (right) for contraversive- and ipsiversive-preferring neurons (mean ± s.e.m across units).

In order to understand the impact of iSPN activation on lSC activity, we recorded single units from the part of SC that received strong ^VLS^SNr projection, in both left and right hemispheres (n=687 units, from 7 mice; Methods) from *Adora2a-Cre* mice expressing CoChR in right striatum iSPN as they performed the lateralized licking task (**Fig. 3e**). We stimulated VLS iSPN as before while recording activity in lSC (**Fig. 1**). We first analyzed lSC activity during trials without stimulation. Activity of individual lSC units displayed selectivity for lick direction that emerged after tone onset and before the choice lick (**Fig. 3e**). Selectivity (the difference in activities in trials cued to preferred vs. non-preferred directions) emerged gradually after tone onset and was maintained before the first lick, indicating lSC has information that could be used to drive the upcoming lick direction (**Fig. 3f**). We further categorized each unit as preferring contraversive vs. ipsiversive licks or as having no directional preference, by comparing the spike counts during the 100 ms window starting at tone onset during correct trials only (see Methods). As a population, units fired more during contra than ipsi trials, with twice as many units that were contraversive lick preferring than were ipsiversive preferring (contra preferring: 296/673, ipsi preferring: 139/673, no preference: 238/673; **Fig. 3g, Extended Data Fig. 5**). Selectivity emerged faster in ipsiversive-preferring units and decayed slowly. However, selectivity gradually increased in contraversive-preferring units, peaking just before first lick (**Fig. 3i, j**). Thus, lSC activity was higher before and during contraversive licking, consistent with lSC involvement in generating contraversive licking (**Fig. 3d**).

### Bilateral push-pull modulation of lSC

We next analyzed trials in which we stimulated iSPN in the right VLS (~25% of trials, **Fig. 4a**). Consistent with the canonical model of the indirect pathway, we observed units in the lSC on the same side (i.e. in the right lSC) that were suppressed by right iSPN stimulation (**Fig. 4b,** top panels). However, we also observed units in the lSC in the opposite hemisphere (i.e. in the left lSC) that were excited by the stimulation **(Fig. 4b,** bottom panels). This was surprising given that the GABAergic ^VLS^SNr cells bilaterally innervate SC (**Fig. 3b**). We also found units on both sides that were not modulated by the stimulation (**Extended Data Fig. 6a**). The overall pattern was consistent across the population with iSPN stimulation generally suppressing activity on the same side (right SC) but exciting activity on the opposite side (left SC) (**Fig. 4c-e**). The effect also persisted well beyond the stimulation window (100ms), suggesting population activity was permanently altered. This effect was also present on both trial types (left-cued and right-cued trials) but with stronger effect on left-cued trials during which behavior was altered by the stimulation and caused ipsilateral bias (**Fig. 1f**).

**Figure 4.**
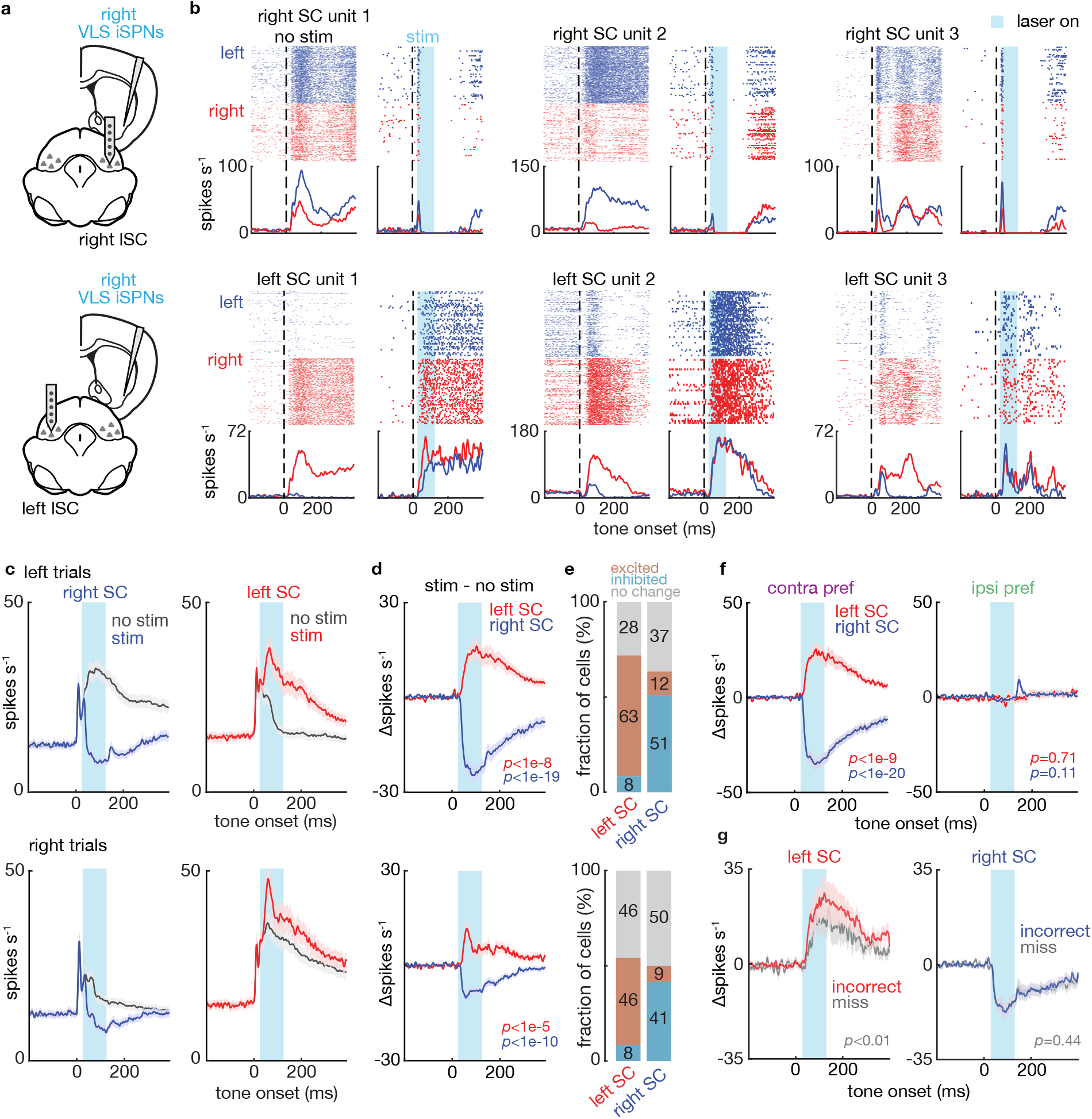
Push-pull modulation of lSC in each hemisphere by iSPN activation. **a.**Schematic of stimulation of iSPNs in the right VLS with simultaneous recording in lSC in the right (top) or left (bottom) hemisphere. **b.** Peri-stimulus histograms of activities for three example units recorded in right SC (top) or left SC (bottom) in left- (blue) and right- (red) cued trials either without (no stim, left panels) or with (stim, right panels) optogenetic stimulation. The timing of the laser is shown in light blue. **c.**Average firing rate of lSC units in the right (blue) and left (red) SC hemisphere during left- (top) and right- (bottom) cued stimulation (blue/red) and no stimulation (black) trials. Only data from neurons with directional selectivity in no stimulation trials are shown (right SC: n = 249, left SC: n= 186). **d.** Changes in firing rate induced by stimulation (Δspikes s^-1^ = activity in stim trials - activity in no stim trials) for neurons in the left (red) and right (blue) lSC. Stimulation increased activity of neurons in the left SC (left trials: P<1e-8; right trials: P<1e-5, two-tailed *t*-test) and decreased activity of neurons in the right SC (left trials: P<1e-19; right trials: P<1e-10, two-tailed *t*-test comparing average activity during the 100 ms optogenetic stimulation). **e.**Fractions of neurons that were significantly modulated. Cells were excited (red), inhibited (blue) or showed no change (grey). All fractions are written for each type. There were more excited vs inhibited units in left SC (left trial: P<1e-26; right trials: P<1e-15; two-tailed binomial test), and more inhibited vs excited units in right SC (left trial: P<1e-10; right trial: P<1e-10; two-tailed binomial test). **f.**As in panel **d** but with data separately shown for contraverisve (left) and ipsiversive (right) preferring units during left-cued trials. Only contraversive preferring neurons were significant modulation by optogenetic (Contra Pref. left SC: n=171, P<1e-9; right SC: n=97, P<1e-20, twotailed *t*-test) (Ipsi Pref. left SC: n=71, P=0.71, right SC: n=70, P=0.11, two-tailed *t*-test). **g.** As in panel d for stimulation trials but including only data from the subset of sessions that had both incorrect and miss outcomes (incorrect: red/blue, miss: grey; Methods) and separating trials based on outcome. Changes in firing rates in left SC neurons were larger during incorrect licking vs miss trials (n=64, P=0.002, two-tailed *t*-test) but did not differ for right SC neurons (n=129, P=0.40, two-tailed *t*-test). All firing rate show mean ± s.e.m. across units.

In order to understand if the net effect of iSPN activity on lSC units depended on their coding properties, we analyzed the effect of iSPN stimulation on direction selective units in right and left lSC separately (**Fig. 4c-d**). Surprisingly, iSPN stimulation specifically modulated contraversive lick-preferring units but not ipsiversive lick-preferring units, in the pattern described above, suggesting the overall pattern of modulation is driven by the effect on contraversive licking-preferring units (**Fig. 4f, Extended Data Fig. 6c-e**).

In a subset of sessions, stimulation caused enough of both incorrect and miss trials to compare the activity in these two kinds of errors. In those sessions, we found that left SC (i.e. contralateral to the stimulation side), but not right SC activity during stimulation predicted behavioral outcome, with higher firing rate during incorrect licking vs miss trials (**Fig. 6g**). In a subset of mice (4/7), we also stimulated in the ITI period, which triggered occasional licks to the ipsilateral side (**Extended Data Fig. 3**). We also observed a similar pattern of inhibition and excitation of lSC hemispheres when the stimulation was applied during the ITI, with activity in left SC differentiating behavioral outcome (**Extended Data Fig. 7**). Overall, the effects of iSPN stimulation on lSC activity depended on location (ipsilateral vs. contralateral side), trial type (right-vs. left-cued trials), and the coding of each neuron (contraversive vs. ipsiversive lick preferring).

### iSPN modulate lSC dynamics along the coding dimension

Covariances in the firing of neurons in a brain region typically makes the neural activity of the population lie in a subspace of vastly lower dimensionality than the number of neurons in the region. We examined if the same is true of lSC activity during the task and if the bases of such a subspace could explain the mouse behavior and effects of iSPN stimulation (**Fig. 5a**). Using only activity measured during control trials (i.e. no optogenetic stimulation), we first computed the coding direction (CD)^24^ with the population activity, defined as the vector along which activity maximally discriminates left-vs right-cued trials. We then applied PCA on the residual subspace orthogonal to the CD to capture the remaining variance (**Fig. 5b**, **Extended Data Fig. 8**). The projection of activity along the CD for left-vs right-cued trials gradually diverged starting at ~35ms after tone onset with maximal separation peaking around 100 ms later (**Fig. 5c, d**). Activity along PC1 increased after tone onset and remained persistently high whereas along PC2 increased sharply and then decayed back to pre-stimulus level after ~80ms (**Fig. 5c**). For these reasons we referred to the former as the ‘persistent mode’ (PC1) and the latter the ‘transient mode’ (PC2). As expected, activity along CD best separated correct left and right trials compared to along the PCs (**Extended Data Fig. 8d**). CD, PC1, and PC2 together captured about 50.4% of the total variance, with CD being the largest contributor (**Extended Data Fig. 8e**).

**Figure 5.**
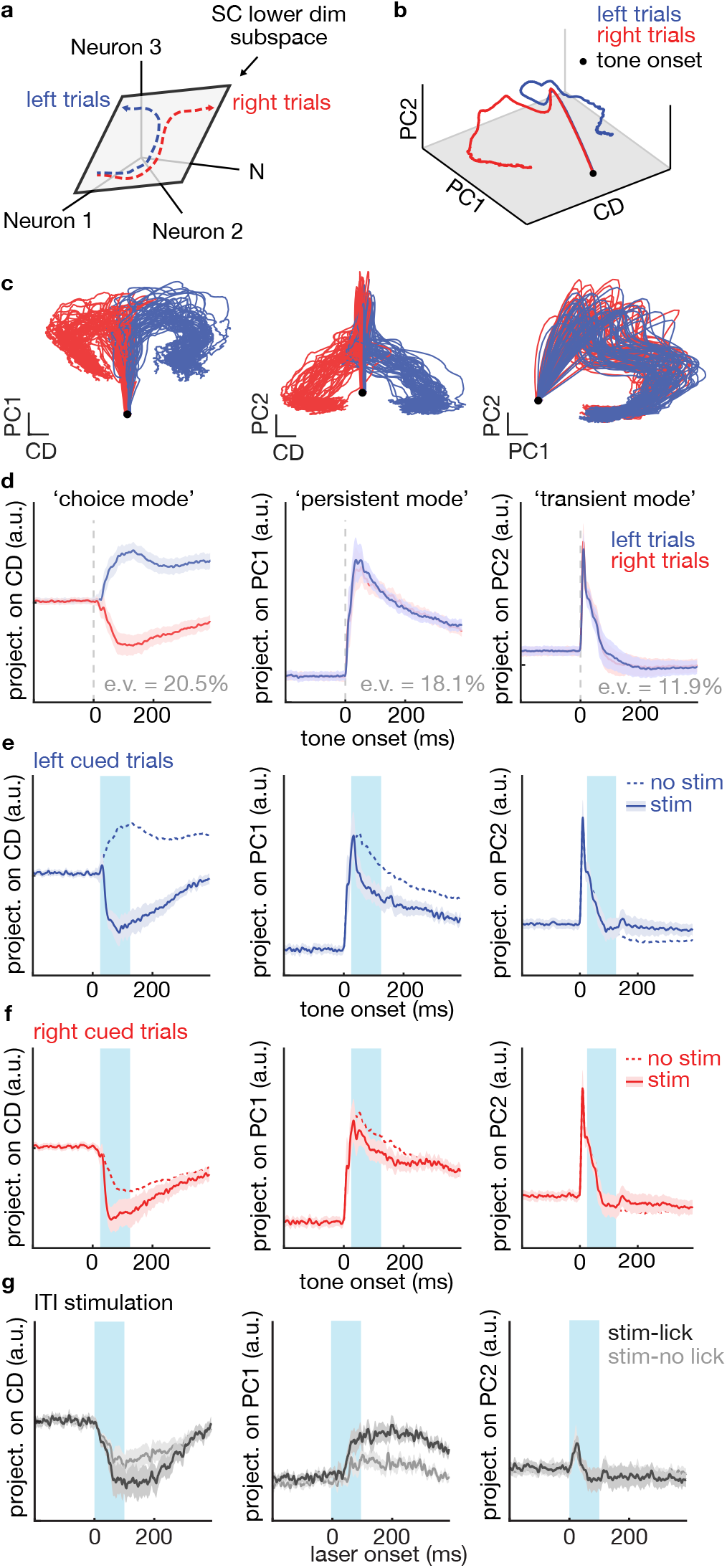
iSPNs activation steers the lSC neural trajectory towards the direction encoding ipsiversive licking. **a.**Schematic showing N-dimensional neural activity in SC is projected onto a lower dimensional subspace (grey) that separates activity during left (blue) and right (red) licking trials. **b.**Mean neural trajectories during left (blue) and right (red) trials projected onto a 3-dimensional subspace (CD, PC1 and PC2; see Methods). Black dot shows the timing of the tone onset (t=0). Only data from no stim trials was used to calculate the subspace. **c.**Neural trajectories plotted in 2-dimensional subspaces defined by combinations of CD, PC1, and PC2. Each line indicates a single bootstrapped trace generated from data during left (blue) or right (red) no stim trials (bootstrapped from units, see Methods). **d.**Activity during no stim trials projected onto CD (‘choice mode’, left), PC1 (‘persistent mode’, middle) and PC2 (‘transient mode’, right) with respect to time. Explained variances (e.v.) along each dimension are indicated. **e-f.** Activities during left-cued (e) and right-cued (f) trials, in stim (thick) and no stim (dotted) trials projected onto the 3 indicated dimensions. Error bars show bootstrapped standard error. g. Effects of stimulation on neural activity during the inter-stimulus interval. Lines show neural activities during the inter-stimulus interval aligned to laser onset for trials in which stimulation did (black) or did not (grey) generate a click. The timing of the laser pulse in shown in light blue in all panels). All error bars indicate bootstrapped s.e.m.

In order to examine if the structure of this subspace was informative for the effects of iSPN stimulation, we projected the activity during optogenetic stimulation trials onto the lower dimensional subspace (i.e. CD, PC1, PC2). Crucially, these trials were not used to define the subspace to compute the CD, PC1 and PC2, and thus allowed us to understand the effect iSPN stimulation in terms of the intrinsic lSC dynamics during control trials. Stimulation in the right VLS pushed the neural trajectory along the CD axis towards the right lick direction, such that during left-cued trials, activity after stimulation looked similar to that in control right-cued trial trajectories (**Fig. 5e-f, Extended Data Fig. 9a**). The effect was smaller during right-cued trials and in other dimensions (**Fig. 5f, Extended Data Fig. 9b**). This indicates that iSPN stimulation pushes the lSC activity specifically along the choice mode.

In order to remove any interaction between the dynamics caused by the cue tone and the optogenetic stimulation, we examined the impact of iSPN stimulation during the ITI in which mice do not normally lick and are not exposed to the tone. Surprisingly, projecting the activity during ITI stimulation along the same axes computed used above (**Fig. 5a-d**) revealed that it recapitulates the neural trajectory observed during right-cued trials (**Fig. 5g**). This was true both during the 100 ms stimulation as well as in the subsequent several hundreds of milliseconds during which the animal normally licks for reward in cued trials. Furthermore, optogenetically-induced activity along PC1 (persistent mode) differed on trials in which the optogenetic stimulation during the ITI did or did not induce a licking bout (**Fig. 5g**, middle panel, **Extended Data Fig. 9c**). Interestingly, small differences of activity along PC1 existed before the stimulation for stimuli that were effective or ineffective at inducing a lick, suggesting that mice were in a different state (**Extended Data Fig. 9d**). Overall, iSPN activity specifically modulates lSC dynamics along the coding direction, even outside the decision window, indicating that it induces ipsiversive licking by steering the neural trajectory in lSC towards the ipsi direction.

## Discussion

Here, we measured behavioral and neural responses after stimulating iSPN in a lateralized licking task. We found that stimulating iSPN suppressed contraversive movement while promoting ipsiversive movement. At the circuit level, this could be explained by concurrent suppression and activation of same and opposite lSC hemisphere, respectively. We also found that iSPN modulate lSC activity specifically along a dimension that differentiates lick choice (i.e. left vs right). Our work shows that the context-dependent behavioral phenotypes observed after iSPN stimulation can be explained by the changes in lSC neural dynamics and suggest a new framework to understand the function of the indirect pathway.

Our results and subsequent conclusions are at odds with two key aspects of previous BG models. First, we found that iSPN stimulation in the VLS suppresses contra licking. Given that the dSPN stimulation in the same region induced contra licking^30^, our work does not support a model whereby co-activation of dSPN and iSPN in the same striatal region cooperate to simultaneously activate a target movement and suppress off-target off-target movements^10^. Second, we found that iSPN activity can both suppress and activate distinct lSC hemispheres, suggesting that iSPN function is not restricted to movement suppression per se, but instead can generate movement itself, thus revealing a unexplored pro-kinetic or movement-triggering function of iSPN.

The circuit mechanisms by which iSPN activity suppresses ipsilateral SC and excites contralateral SC or by which it modulates only contraversive-licking preferring neurons are unclear. Given that SNr projects to lSC on both sides, one possibility is that iSPN differentially modulate contra and ipsi lSC by differentially modulating SNr populations that target different SC hemispheres^5,38^. This suggests a specificity of wiring of the iSPN projection through the GPe and STN that has not been explored experimentally. Another possibility is that unilateral lSC inhibition causes disinhibition of contra lSC through long range and net inhibitory inter-collicular projections^39,40^. Further work will be needed to tease apart these different models of contralateral lSC excitation.

SC has been extensively studied in the context of action and target selection^31,32,34,35,41^. Differential activity between SC hemispheres correlates with choice accuracy^34^, reaction time^42^ and perceptual judgements^43^. One popular model posits that this differential activity, or winnertake-all dynamics can emerge via an attractor network implemented via inhibition between SC hemispheres^35,44,45^. Here, we show that iSPN activation is sufficient to generate this differential activity (**Fig. 4**) which, in our task, corresponded to pushing activity along the CD direction towards the neural activity subspace associated with ipsiversive movements (**Fig. 5**). Thus, one way to conceptualize the effect of iSPN activity is that it pushes the neural trajectory away from the contraversive movement side along a decision axis. Depending on the other motor programs that feed into SC by other circuits outside the BG, this iSPN induced neural trajectory might translate into movement suppression (fall back into resting state) or ipsiversive movement generation (push towards ipsi side) (**Fig. 2, Extended Data Fig. 2, 3**).

While we studied iSPN in the context of a lateralized behavior, it is possible that a similar mechanism might exist for pairs of categorically different actions (e.g. licking vs locomotion). A previous study that stimulated iSPN during a lever press task found that the stimulation aborted lever pressing behavior, while inducing locomotion in the chamber^11^. Testing this hypothesis fully would require activating different ensemble of iSPN in the same striatal hemisphere encoding distinct actions, while monitoring SC activity. Similarly, we studied a learned action motivated by a water reward and it is unknown if similar mechanisms influence choices between spontaneous actions or action sequences in the absence of overt motivation by immediate reward. Lastly, our focus is on a functionally and anatomically defined pathway originating in the lick-associated VLS and terminating in the lSC. Further studies are needed to preform similar analyses in other anatomically or functionally defined striatal subregions.

Although our work did not address what iSPN activity encodes, previous studies have shown that iSPN are more active after mice experience a negative outcome^46–48^. Thus, the function of the iSPN might be to associate a negative outcome with a specific action in order to avoid them in the future. In this model, the ‘negative reward prediction error’ signaled by dips in dopamine neuron activity^49,50^ would strengthen inputs onto iSPN^51^ such that the subsequent increased iSPN activity can more effectively suppress the corresponding action and promotes alternative options. Our findings suggest that iSPN have the capacity to implement this computation, which would be beneficial for the survival of an organism.

## Acknowledgements

We thank members of the Sabatini laboratory and W. Regehr, M. Andermann, N. Uchida S. Gershman for helpful discussions. We thank J. Levasseur for mouse husbandry and genotyping, and J. Saulnier and L. Worth for laboratory administration. We thank W. Kuwamoto, J. Grande, M. Ambrosino, B. Pryor and E. Lubbers for assistance with behavioral experiments and histology. This work was supported by the NIH (grant no. NINDS R01NS103226), a P30 Core Center Grant (grant no. NINDS NS072030), an Iljou Foundation scholarship and a grant from the Simons Collaborative on the Global Brain.

## Author Contributions

J.L. and B.L.S. conceptualized the study, wrote the original draft, and reviewed and edited the manuscript. J.L. performed experiments and analyzed the data. B.L.S. supervised the study and was responsible for acquisition of funding.

## Declaration of Interests

Dr. Sabatini is a founder and holds private equity in Optogenix. Tapered fibers are commercially available from Optogenix were used as tools in the research.

## Methods

### Mice

All mouse handling and manipulations were performed in accordance with protocols approved by the Harvard Standing Committee on Animal Care, following guidelines described in the US National Institutes of Health Guide for the Care and Use of Laboratory Animals. For behavioral experiments, we used male and female (3~6 months old) *Adora2a-Cre* (B6.FVB(Cg)-Tg(Adora2a-cre)KG139Gsat/Mmucd, 036158-UCD) from C57BL/6J backgrounds acquired from MMRRC UC Davis. For muscimol infusion experiments, we used wild type (C57BL/6NCrl, Charles River) mice (age >P75) were used.

### Surgery and viral injection

All mice underwent headpost/fiber surgery before training, and craniotomy surgery after training, prior to electrophysiology. This minimized the duration of brain surface being exposed. Mice were anaesthetized with isoflurane (2.5% in 80% oxygen). Using a stereotaxic frame (David Kopf Instruments, model 1900), mouse’s skull was exposed and leveled (David Kopf Instruments, Model 1905). The patch of skin covering the skull was cut and removed. ~300 μm diameter craniotomy was made with a drill (David Kopf Instruments, Model 1911) for each viral injection. Viruses were injected using a pulled glass pipette (Drummond Scientific Company pipettes) that was cut at beveled (~ 30 degrees, 35 ~ 50um inner diameter), and a syringe pump (Harvard Apparatus, #84850). Viruses were frontloaded at a rate of 500nl/min, then the pipette slowly lowered into the target region. The pipette was first lowered 300 μm deeper than the target dorsoventral coordinates. The pipette was left in the brain for 5 minutes before injection began, at a rate of 75 nl/min. After infusion, the pipette was left in place for a 5 minutes before it was slowly withdrawn. For fiber implants, a stereotaxic cannula holder (SCH_1.25, Doric) was used to hold the fiber and slowly lower the fiber into the brain. The fiber and headpost were secured on the skull using loctite gel (McMaster-Carr 74765A65) and zip kicker (Pacer Technology). A wall surrounding the site of recording was made using loctite to contain the saline bath for electrophysiology grounding. The site of recording was marked, and the wall was filled with silicone gel (Kwik-Sil, World Precision Instruments). Mice were given pre- and post-operative oral carprofen (MediGel CPF, 5 mg/kg/day) as an analgesic, and monitored for at least 5 days. For craniotomy surgery, the silicon was removed and a craniotomy was made by drilling the skull with a 340 um diameter drill bit. The craniotomy was extended 300um medial/lateral and anterior posterior. Care was taken not to damage or puncture dura, as it resulted in more infection of the craniotomy.

Viral injection for striatal tapered fibers photostimulation was done in a similar way as previously described^30^. Briefly, AAV2/9-hSyn-FLEX-CoChR-GFP (UNC vector core) was injected in the medial and lateral part of striatum (titer: 5×10^12^ gc/ml, injection volume: 300 nl per site, total 1200 nl). Lateral striatum virus injection and fiber implant was done at an angle. All coordinates were as follows (AP/ML/DV relative to bregma and dura, in mm): DMSVMS: 0.5/1.25/-3.25 and 2.15; DLSVLS: 0.5/3.4/-3.35 and −2.15, at 14.5 degrees; lateral SC: −3.4/1.5/variable).

### Histology and immunohistochemistry

Mice were euthanized and perfused transcardially with 1M PBS followed by 4% PFA (1M). After 24 hours post-fix in 4% PFA, brains were equilibrated in 30% sucrose solution until they sank at the bottom. Brains were then sliced (50um thick) using a cryostat. Slices were mounted on slide glasses with DAPI mounting medium (VECTASHIELD, H-1200) and imaged under a widefield microscope with a 10x objective (VS120 OLYMPUS). In some mice, for localizing the location of the tapered fibers, we immuno-stained for glial fibrillary acidic protein or GFAP (Agilent Technologies, Z033429-2, 1:500 dilution ratio).

### Behavior

We designed a lateralized licking task in which mice had select between two lateralized actions and report their decision by licking the relevant spout instructed by the tone frequency. Mice were headfixed and placed inside a plastic tube^52^. Each trial began by an intertrial-interval (ITI) during which mice were required to withhold licking. ITIs were chosen randomly between 2000 ~ 4000 ms. Any lick during the ITI reset the clock, but did not change the ITI duration. If no licks were detected during the ITI, a 50 ms duration tone of either low (tone A, 3kHz) or high (tone B, 12kHz) frequency was played. Mice had to lick the left spout (tone A) or right spout (tone B), after which a small water drop (1 ~ 1.4ul) was immediately dispensed from the corresponding spout. Not licking within a response window (500 ms relative to tone onset) resulted in a miss trial, and a time out period (6000 ms). Each session lasted for an hour or until the mouse had multiple consecutive miss trails (~10 miss trials).

Mice were trained in a series of stages in order to reach final expert performance. Mice were first water deprived (up to 85% baseline weight) and habituated to head fixation. This was followed by water delivery on the rig via one of the spout centered in front of their mouth. A dummy version of the final task was used in which only one tone type was played, wither shorter ITI (1000 ~ 2000 ms), longer response window (1000 ms) and shorter time out period (1000 ms). This version of the task was meant to teach the mouse to associated the tone with licking. A small water drop was manually dispensed initially when the tone was played to help the training. Mice were then trained to lick sideways by positioning one of the side spout at the center initially and gradually moving it to the final position. Mice learned to track the position of the spout and lick sideways within 1 ~ 3 days. The same procedure was repeated for both sides, at least two times for each side (each spout ~60 rewards delivery per repetition). Mice were then trained on the final task. Duration of ITI, response window and time out period was gradually adjusted to the final values. During the training phase, we repeated the same tone after an incorrect trial, in order to prevent the mice from learning a strategy of licking only one spout and still collecting rewards.

Mice were trained for at least two weeks (from habituation), after which they were trained until they reached 85% correct performance. For extinction (Fig. 2), mice underwent the same training procedure, after which the right spout devalued by not dispensing any water reward even after a correct lick. During extinction, we removed timeout period given that mice learned to not lick to the devalued spout (Fig. 2e). We also trained mice to lick only one spout (Extended Data Fig. 2). Mice underwent the same procedure but was only ever exposed to the left spout throughout training. The right spout was still present and available to the mouse.

### Behavioral set-up

Behavioral data was acquired and saved using Arduino (MEGA 2560) and CoolTerm. Licks were detected by recording the voltage drop between the spout and the tube, similar to previously described (Ref) The inside of the tube was taped with copper foil and grounded. Solenoids (The Lee Company, part number LHQA0531220H) were connected to a 20ml syringes, acting as water reservoirs, and opened for a short duration to deliver water rewards. Water reward size was calibrated via adjusting solenoids opening time (~20 ms). Water delivery spouts were made using blunt syringes needles (18 gauges). They were glued in parallel, separated by 6.5mm, and connected to the solenoids via tubing (Cole-Parmer, # EW-06460-34). A speaker (Madiasound, parts number: tw025a20) connected to an amplifier (FOSTEX, parts number: AP05) was positioned underneath the tube, connected to the Arduino to deliver tones during the task.

### Photostimulation

We photoactivated iSPNs in striatum by expressing CoChR in striatum and delivering blue light (473nm, Optoengine). For functional mapping, two tapered fibers (0.66 NA, emitting length 2 mm, implant length 2.5 mm, Optogenix) were implanted to stimulate a total 8 striatal sites using an optical setup used previously. Only one fiber was stimulated per session (4 sites per session). Stimulation was randomly interleaved, and was deployed 20 ~ 30% of the time, to minimize persistent behavioral phenotype due to repeated stimulation. We did not observe any gross persistent effect on baseline performance across session due to stimulation. Each photostimulation session consisted of stimulation trials on both left and right trials, across 4 striatal sites from one fiber. We ran two sessions per fiber per mouse. For other experiments (extinction, one spout training and extracellular recording), only one fiber was implanted per hemisphere, and only one site (VLS) was targeted for photostimulation. We calibrated the power level for each depth by adjusting the power at the end of the patchcord (before fiber entry) to be 100 uw. Stimulation consisted of a constant 100 ms pulse, delivered 25 ms after tone onset. In order to stimulate distinct depths along the tapered fiber, we used an optical system for delivering different modes of light onto the back of a high NA patchcord (0.66 NA) connecting the tapered fiber, similar to the work previously described^30^. A custom code in Matlab was used to control the optical set-up via a data acquisition interface (National Instrument) and communicate with the Arduino.

### Electrophysiology

We performed *in-vivo* extracellular recoding in lateral superior colliculus while mice were performing the licking task. Mice underwent viral and fiber implant surgery, after which they were trained on the main task for two weeks. Recording mice only received a single fiber in the lateral part, targeting VLS. iSPNs in VLS were stimulated to characterize the behavioral phenotype. All recorded mice show similar behavioral phenotype as reported in previous experiments where only stimulation was performed. One day before the first recording session, one small craniotomy above each lateral SC were made (~0.5 mm in diameter). Care was taken to not remove the dura when drilling through the skull. The craniotomies were covered with silicon gel (Kwik-Sil, World Precision Instruments). During subsequent recording sessions, the silicon gel was removed and the craniotomy was filled with clean saline solution. 64 channel silicon probes (A2×32-5mm-25-200-177 or A4×16-Poly2-5mm-23s-200-177, NeuroNexus Technologies) were lowered slowly in brain until it reached the target depth (lSC: −2.2 ~ −2.6 mm relative to dura). We explored different location within the craniotomy in order to record from diverse locations within lSC. We dispensed water rewards while lowering the probe and looked for signals locked to rhythmic licking. Cells within a narrow layer spanning about 400um around lSC consistently fired in relation to licking. Once reaching the target depth, we left the probes for an additional 5 minutes for the surrounding tissue stabilize before starting the recording session. For most mice, only a single depth along the tapered fiber (VLS) was stimulated while recording.

We alternated the recoding location of lateral SC (ipsi or contra relative to fiber location) every day and later analyzed lSC units separately based on recording location (Fig. 4). We performed recording until the craniotomies became too unhealthy to record from, performance degraded due to repeated insertion of the silicon probe, or the number of observable units in lSC dramatically decreased (range: over 2 weeks). During the last session, we marked the center of the craniotomy with probes coated with Dil and later confirmed the recording location in histological slices.

Signals were acquired through OmniPlex Neural Recording Data Acquisition System (Plexon Inc). Signals from each channel were filtered (analog filter 0.1-7500 Hz; digital filter 0.77Hz Highpass), digitized at 40kHz, and single units were manually sorted using Offline Sorter (v3.3.5, Plexon Inc). Units were first detected using a hard threshold (below −44.63uV). Neighboring channels were grouped into tetrodes to aid sorting. Principal Component feature space was visually inspected and used to manually draw boundaries of each putative single unit cluster. Artifacts due to spout contact were clearly visible in all channels and easily removed using non-linear energy/energy dimension.

### Muscimol infusion

We infused muscimol (Sigma-Aldrich) in lateral SC unilaterally while mice were performing the lateralized licking task. A craniotomy (~0.6mm diameter) was made above each lateral SC one day prior to muscimol infusion. We used a glass pipette frontloaded with muscimol via a syringe pump (see Surgery and viral injection). While the mouse headfixed, the silicon gel above the craniotomy was removed and the injection pipette was slowly lowered into the target depth (2200um below dura). The pipette was left in the brain for 5 minutes before starting the behavioral session. When the mouse had performed 200 ~ 250 trials, we started infusing muscimol (450-500ng/ul)^37^ at a rate of 50nl/min and a total volume of 100-150nl. All mice displayed licking deficit within the first 5 minutes of the start of infusion. We compared the performance pre and post muscimol infusion (Fig. 5c). For post muscimol infusion trials, we analyzed the last 200 trials of the session in order to take into account the time for muscimol to diffuse in the tissue. After 1 hour, we aborted the session, the pipette was slowly raised, and the craniotomy was covered with fresh silicon gel. Some mice received a mixture of muscimol and Cholera Toxin Subunit B (Recombinant) Alexa Fluor (647 conjugate, Thermo Scientific) to localize site of infusion. All mice fully recovered from previous muscimol infusion session and no performance deficit was observed on the next session, during pre-infusion control trials. At one infusion site, we noticed that baseline performance was low from the beginning, possibly due to damage from the infusion pipette being lowered into the brain. Each mouse received two infusion sessions (one per site), and all sessions were combined for the analysis (Fig. 3c).

### Behavioral data analyses

We categorized each trial outcome as correct, incorrect or miss. Correct trials were trials in which mice liked the correct spout (tone A-left, tone B-right) within a response window (500ms). Incorrect trials were trials in which mice licked the wrong spout. Miss trials were trials in which mice did not initiate any licks within the response window. We quantified mouse’s performance by counting the fraction of correct, incorrect and miss trials for a given session (Fig. 1).

Functional map of striatal site effective at changing behavior was determined using hierarchical bootstrapping to account for variability across mice, sessions and trials (Fig. 1e). We tested against the null hypothesis that the stimulation did not change the fraction of correct/incorrect/miss trials. In each round of bootstrapping, we re-sampled data by separately replacing from mice, sessions within each mouse, and trials (both stim and no trials shuffled) within each session. We then computed the performance change on the re-sampled data set. Bootstrapping 10^5^ times produced a distribution of performance changes that reflected the behavioral variability, and the one-tailed p-value was the fraction of times in which bootstrapped data produced equal or greater change in performance then that observed. To compare performance changes after VLS iSPNs stimulation, we performed *t*-test (two-tailed) on the percentage outcome for stim and no stim trials across all sessions (Fig. 1f, Fig. 2b, d, Extended Data Fig. 1, 2). For extinction experiment, for each mouse, we quantified the fraction of correct/incorrect/miss trials during stim/no stim trials for left and right trials, and before/after extinction (Fig. 2). We ran at least two stimulation session for each condition (pre vs post extinction) and averaged the percentage outcome across sessions. In order to test if extinction changed the probability of licking incorrectly, we first computed the change in incorrect rate after stimulation (Δincorrect) to account for baseline incorrect rate and compared it before and after extinction (paired t-test, one-tailed, Fig 2b, d).

### Electrophysiological data analyses

We collected extracellular recording data from 7 mice, 71 individual session, comprising of 617 units (left SC/right SC=294/379). Average number of sessions per mouse was 10.3 sessions (range: 7-17). Average number of units per session was 9.3 units (range: 1-23). Average number of units per mouse was 90.1 units (range: 57-203). All units were pooled together for analysis. We smoothed the firing rate traces with a gaussian window for display purposes (Individual units: 20ms window, mean across units: 10ms window).

For each unit, we determined its coding preference (contra preferring vs ipsi preferring) by comparing the spike count in the first 100ms window after tone onset during left vs right correct trials (two-tailed *t*-test, p<0.05). Each unit was categorized into contra preferring, ipsi preferring, or no preference if it did not pass the p-value threshold. Contra and ipsi preferring units were termed ‘selective units’. To compute selectivity, for each unit we computed the mean firing rate during preferred minus anti preferred trial type. Selectivity was then averaged across units to give a measure of population selectivity (Fig. 3f). In order to test for bias in the population selectivity, we compared spikes count during ipsi vs contra trials at various windows after tone onset (Fig. 3g, Extended Data Fig. 5f). Coding preference could reflect distinct cell type within SC (e.g. excitatory vs inhibitory). However, we did not observe any difference in spike waveform features, although contra preferring units tended to have higher mean firing rate (Extended Data Fig. 5c-e).

To quantify the effect of stimulation on lSC activity, we grouped all units recorded on the left SC or right SC and computed the change in firing rate after stimulation (Δspikes s^-1^=no stim spikes s^-1^-stim spikes s^-1^, Fig. 4d). We also quantified whether the effect of stimulation was significant by comparing the spike counts during the stimulation window (100ms) vs control window (25 ~ 125ms relative to tone onset). Each unit was categorized into significantly excited, inhibited, or no change (Fig. 4e). The above analyses were done for all selective units (Fig. 4c-e, Extended Data Fig. 6b, f), contra preferring units only (Fig. 4f, Extended Data Fig. 6c), ipsi preferring units only (Fig. 4f, Extended Data Fig. 6d, f), or units with no preference (Extended Data Fig. 6e, f).

In order to test whether the activity after iSPNs stimulation could predict behavioral outcome, we analyzed a subset of sessions during we had more 5 trials for each outcome (miss and incorrect). A total of 29 sessions, 193 units were analyzed (Fig. 4g). We compared the spike counts during the stimulation window (100ms) vs control window (25 ~ 125ms relative to tone onset).

In a subset of mice (4/7), we stimulated iSPNs during ITI period. This allowed us to test the effect of stimulation on lSC activity at resting state, without the interaction with the tone. We conducted similar analysis as above, to test whether the lSC activity after iSPNs stimulation could predict behavioral outcome, by analyzing the spikes count during the stimulation window (Extended Data Fig. 7).

### Dimensionality reduction

We applied dimensionality reduction technique similar to previous studies in order to understand the effect of stimulation on lSC activity^24,53^. We assumed all n units from different sessions/mice could have been recorded simultaneously and were pooled together to make a trial-averaged matrix **x** (n × t dimensions) aligned to 1^st^ lick, with each row representing a single unit, and each column representing a single time bin. We found an n × 1 vector, in the n dimensional activity space that maximally separated the response vectors in correct lick left trials (**x**(t)_left_) and correct lick right trials (**x**(t)_right_), termed the coding direction (**CD**). **CD** was computed by subtracting the activity of left – right trials during a 200ms window centered around the time of 1^st^ lick (−100 ~ +100ms relative to spout contact), and divided by it’s length, giving a unit vector **CD**. Projecting activity along **CD** (**CD**^T^**x**) allowed us to separate trajectory for left vs right trials (Fig. 5d, Extended Data Fig. 8d). By construction, **CD**^T^**x** was positive during left trials. Although **CD** was computed using the time of 1^st^ lick, we used activity **x** aligned to tone onset for computing projections, to make comparison between stimulation and no-stimulation trials easier. Separability for trial type was defined as the difference between left vs right trials projection along a specific dimension (e.g. **CD**). We also explored different time windows for the choice of **CD** (0~100ms relative to tone onset, −100~0ms relative to spout contact), and obtained similar separability for left vs right trials (Extended Data Fig. 8d).

In order to capture the remaining variance in the data, we built a matrix consisting of trial-averaged activity for n units during left and right trials, with t time bins. Left and right trials were concatenated, giving a n × 2t matrix. We then removed the component along **CD** by subtracting the projection along **CD** giving **x**_⊥CD_ = **x** – (**CD**)(**CD**^T^**x**), which is the subspace orthogonal to **CD**. We then applied standard PCA to this **x**_⊥CD_, giving PCs that capture variance orthogonal to **CD**. We used time points −400 ~ +400ms relative to tone onset for PCA. Data was centered, but not normalized, thus preserving differences in firing rate across units. Only correct control trials were used to compute the **CD** and **PC**s. We also tried to use PCA without computing and subtracting **CD**. This gave similar results as the approach mentioned above, with **PC2** separating left vs right correct trials (Extended Data Fig. 8c, d).

In order to understand the effect of iSPNs stimulation on lSC activity, we projected stimulation trials activity matrix onto different dimensions (**CD**, **PC1**~**PC5**). Importantly, stimulation trials were not used to compute different dimensions. To quantify the magnitude of stimulation along specific dimensions, we computed the difference between stimulation and no stimulation trial projection along specific dimensions (Extended Data Fig. 9b). We took all correct/incorrect/miss trials for no stimulation trials, in order to make stim vs no stim comparison fair. For stimulation during the ITI, we similarly projected the ITI stimulation trials activity matrix onto different dimensions. Given that only a subset of mice were given ITI stimulation, we re-calculated **CD** and **PC**s for that subset of data (Fig. 5g)

All error bars for projections along lower dimensional space were computed using bootstrapping across units. Every bootstrap consisted of resampling units with replacement and computing **CD** and **PC**s *de novo* (5000 times). P-values were the fraction of times a bootstrap resulted in the opposite sign of that experimentally obtained. PCA results after bootstrapping can be unstable (sign flipping) due to the indeterminacy of the sign of PCA loadings. We used an approach previously described^54^ to assign a sign to **PC**s that most resembles the direction of the data after each bootstrap. For each **PC**s, we changed the sign of the **PC**s so that the sum over the dot product of the **PC** and data points would be greater than zero: 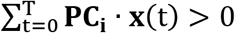. For each **PC**s, we explored and chose different timepoints that did not result sign flipping.

### Statistical analyses

All statistical analyses were performed using custom code written in Matlab (MathWorks, Natick, MA, USA). We used two-tailed *t*-test for all statistical comparisons unless stated otherwise. For functional mapping of striatum (Fig. 1e), analysis of low dimensional subspace (Fig. 5), we used bootstrap (see **Behavioral data analyses** and **Dimensionality reduction**). The significance level was not corrected for multiple comparisons. No statistical methods were used to predetermine sample sizes, but our sample sizes are similar to those reported in previous publications. Data distribution was assumed to be normal, but this was not formally tested.

### Data and code availability

The data that support the findings of this study are available from the corresponding author upon reasonable request. The code used for analysis (Matlab) is also available from the corresponding author upon reasonable request.

---

**Extended Data Fig. 1.**
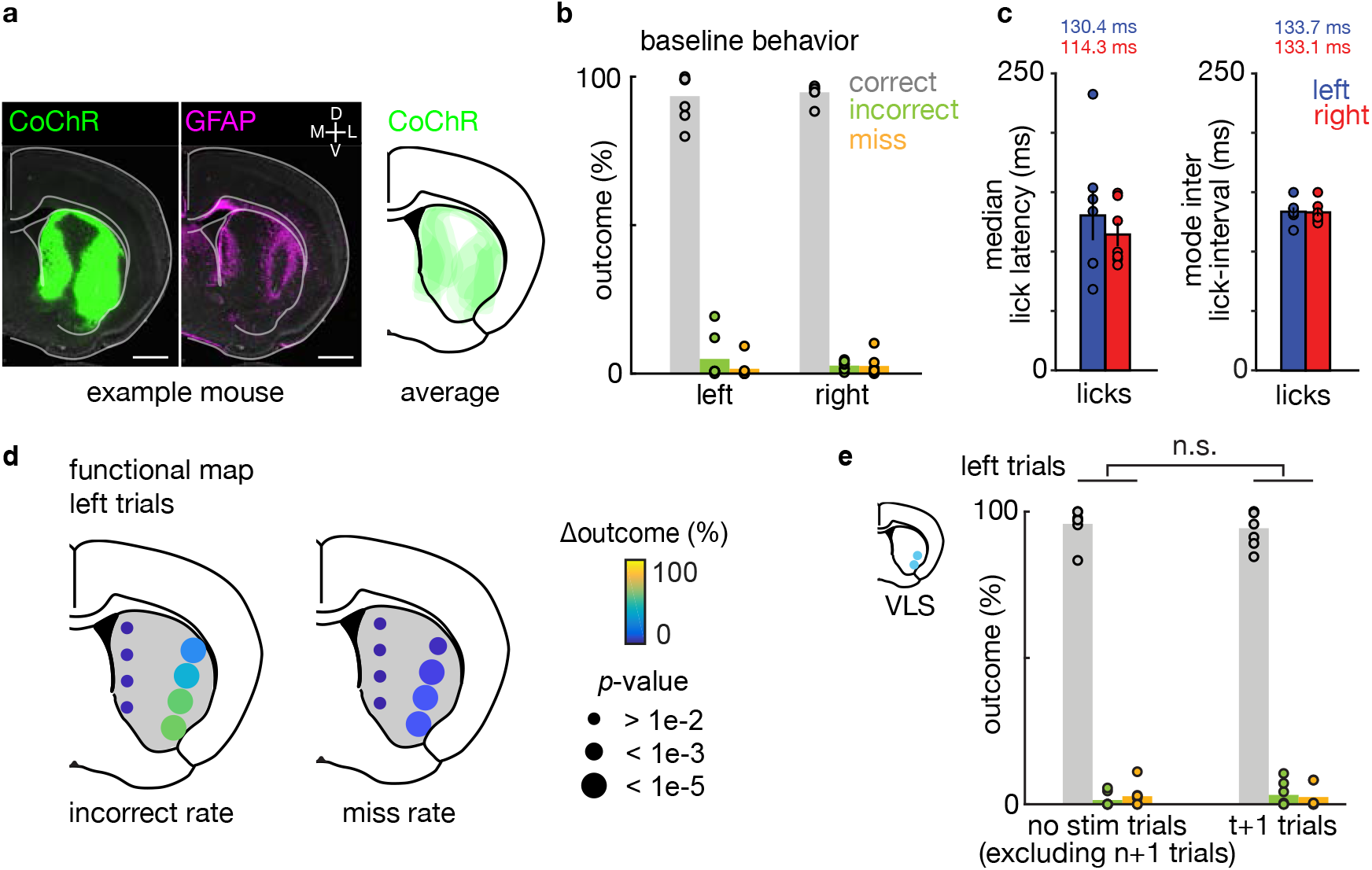
Histology, baseline behavior, and effects of iSPN stimulation on the next trial. **a.** Example mouse histology *(left)* showing CoChR expression (green) and the tapered fiber location as revealed by glial fibrillary acidic protein (GFAP) staining (magenta). Scale bar (1 mm). The average CoChR expression in striatum averaged across mice is also shown *(right).* **b.** Baseline behavior after two weeks of training. Percentages of correct (grey), incorrect (green) and miss (orange) outcomes for left- and right-cued trials (n=7 mice). **c.** *left,* Median lick latency measured from tone onset to spout contact for left- (blue) and right- (red) cued trials. *right,* Mode inter-trial-interval for licks to the left (blue) and right (red) ports. **d.** Functional map of optogenetic perturbations at 8 striatal sties showing changes in percentages of incorrect (left) and miss (right) outcomes (see Fig. 1e). The color and size of each circle denote the effect size and p-value (bootstrap), respectively (n=5 mice, 9 sessions). **e.** Effect of VLS iSPNs stimulation on the next trial (n+1 trial) relative to control trials (excluding all n+1 trials). For n+1 trials, only those following left-cued trials were included as optogenetic stimulation only affected left-cued trials (i.e. contraversive to the stimulation site in the right striatum) (n=7 mice) (see Fig. 1e; Methods) (n.s.: P>0.05, two-tailed *t*-test).

**Extended Data Fig. 2.**
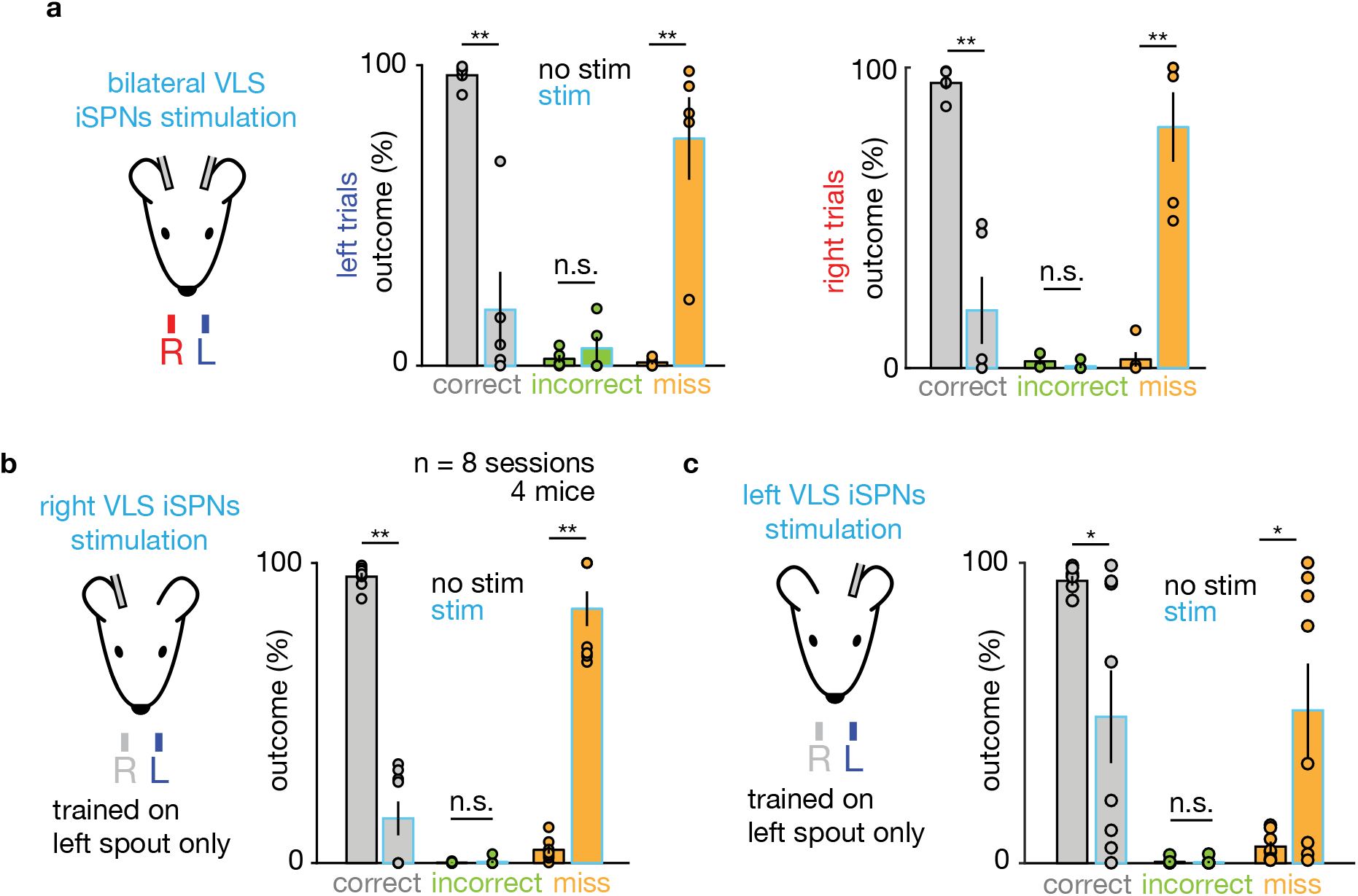
The effects of bilateral iSPNs stimulation on mice trained on two spouts, and of unilateral stimulation on mice trained on only one spout. **a.** Summary plots for the outcomes for no stim (black) and stim (light blue) trials, during left- (center) and right- (right) cued trials (n = 5 mice). Optogenetic stimulation significantly decreased the correct outcome rate and increased the miss outcome rate but did not change the incorrect outcome rate (**P < 0.001, two-tailed *t*-test; n.s.: P > 0.05). **b.** Summary plots for effects of right VLS iSPNs stimulation in mice only trained to lick the left spout (see Methods). Stimulation decreased correct outcome rate and increased the miss outcome rate, but failed to increase incorrect outcome rate (i.e. the rate of licking to the right spout which the mice were never trained to lick) (**P < 1 × 10-8, two-tailed *t*-test; n.s.: P > 0.05). **c.** As in panel **b** for left VLS iSPNs stimulation (*P < 0.05, two-tailed *t*-test; n.s.: P > 0.05).

**Extended Data Fig. 3.**
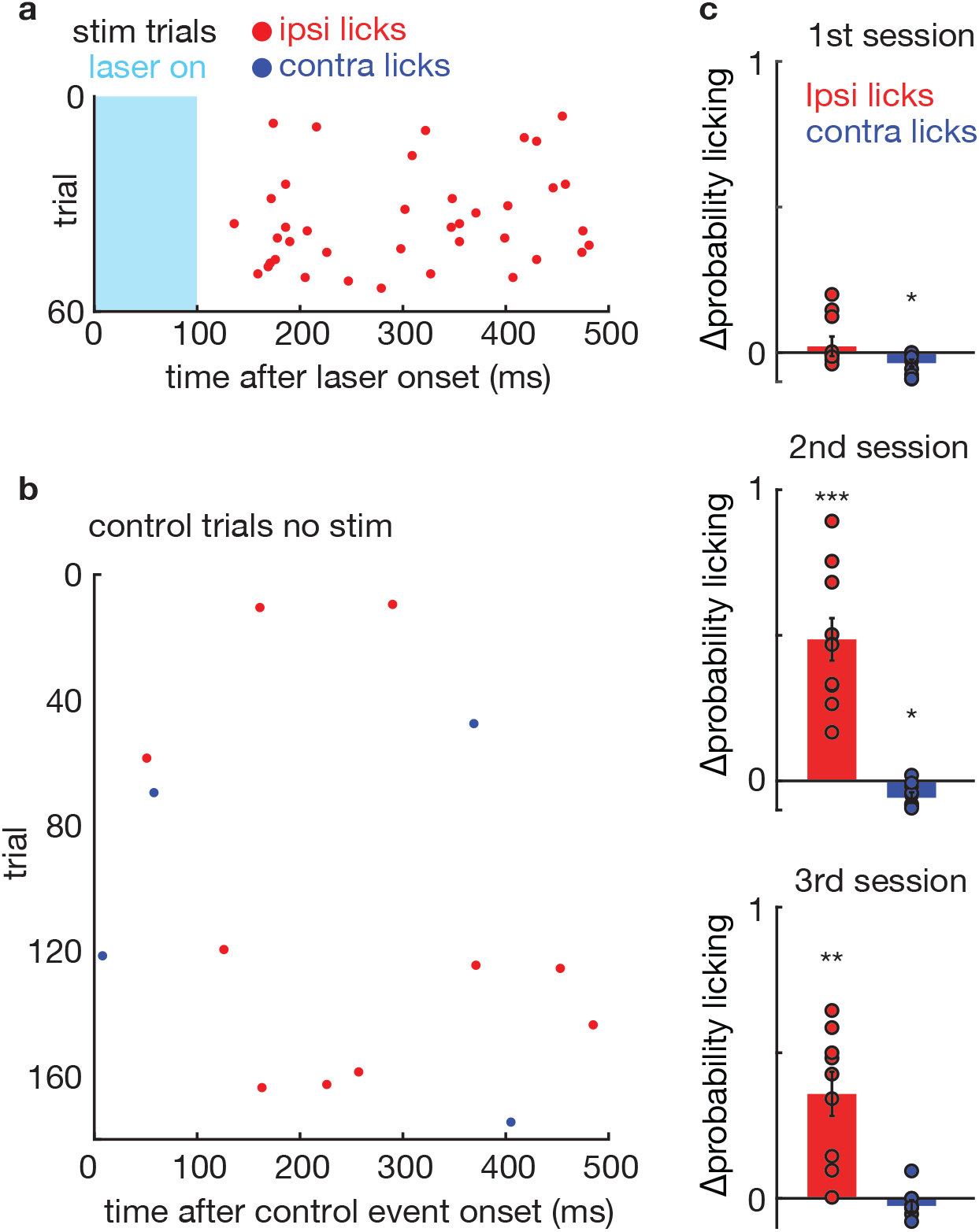
Repeated iSPNs stimulation during ITI triggers ipsiversive licking. **a-b.** Rasters showing the timing of licks in a session during which stimulation during the intertrial interval (ITI) successfully caused ipsilateral licking. Panel **a** shows all the stimulation trials with each lick color-coded by the lick direction (ipsi-red, contra-blue) and are time-aligned to laser onset. Panel **b** is similar but shows licks aligned a control pseudo-event generated by the behavioral control software in which stimulation could have happened but was not delivered. Therefore, this shows the baseline licking rate during the ITI period. **c.** Change in probability of licking after optogenetic stimulation during the ITI relative to control trials (as in panel b) (n = 10 mice for 1^st^ and 2^nd^ session, n = 9 mice for 3^rd^ session). Stimulation caused ipsilateral licking from 2^nd^ session onwards, and weakly suppressed contralateral licking relative to baseline (***P < 1×10^-4^, **P < 0.005, *P < 0.05).

**Extended Data Fig. 4.**
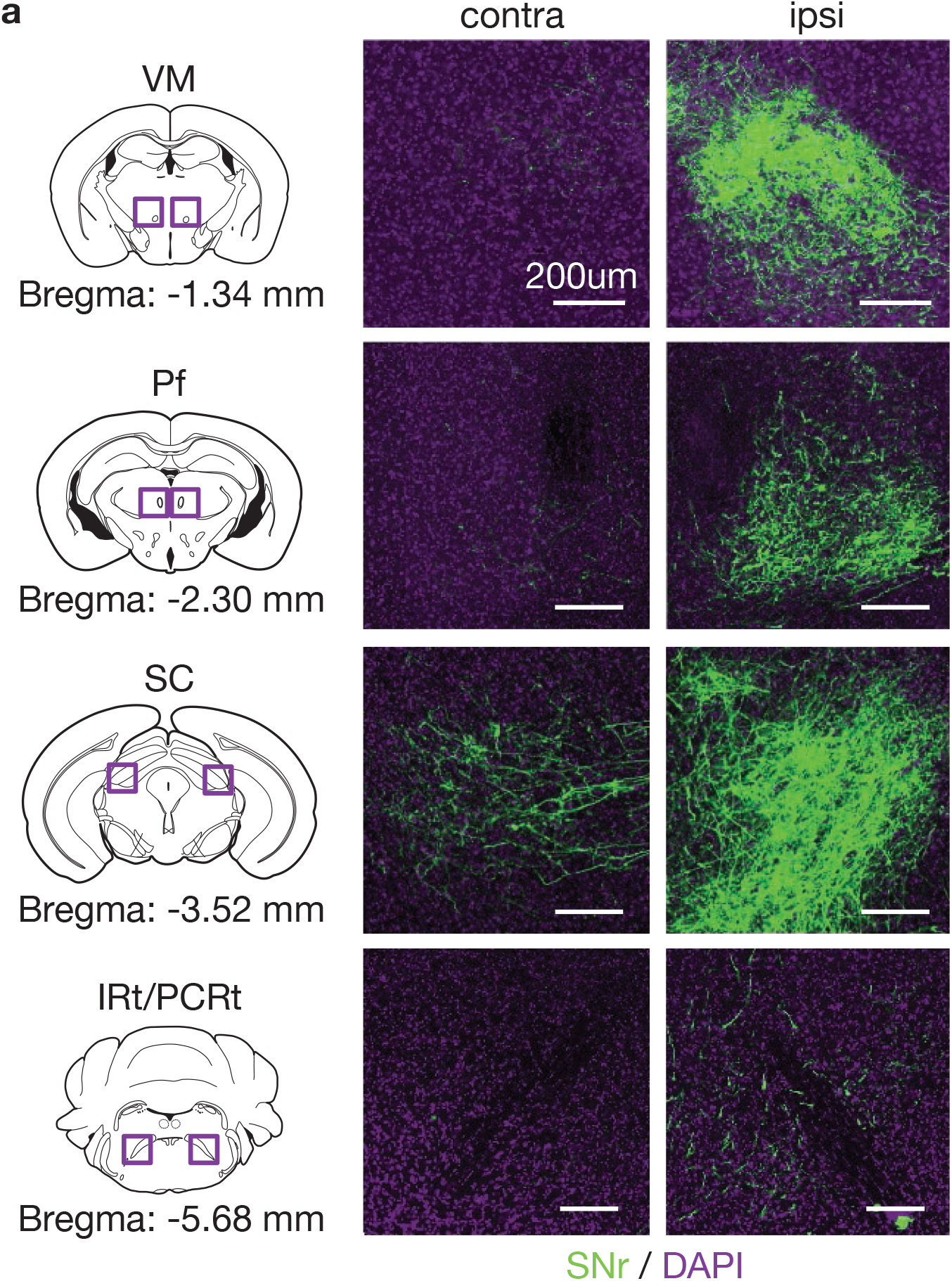
VLS recipient SNr projects to the contra side in SC. *left,* Schematics of coronal sections and coordinates relative to bregma. VM: ventromedial thalamus; Pf: parafasciular nucleus; SC: superior colliculus; IRt/PCRt: intermediate reticular formation/parvocellular reticular formation. *right,* histological examples showing SNr axons (green) labelled via anterograde tracing (see Main text, Fig. 3b) and DAPI (purple). The left and right columns show contralateral and ipsilateral sides, respectively, relative to the labeled SNr cell bodies (i.e. the injection side). Midline crossing SNr axons were only seen in lateral SC. Similar results were observed in total of n=3 mice. Scale bars: 200 μm.

**Extended Data Fig. 5.**
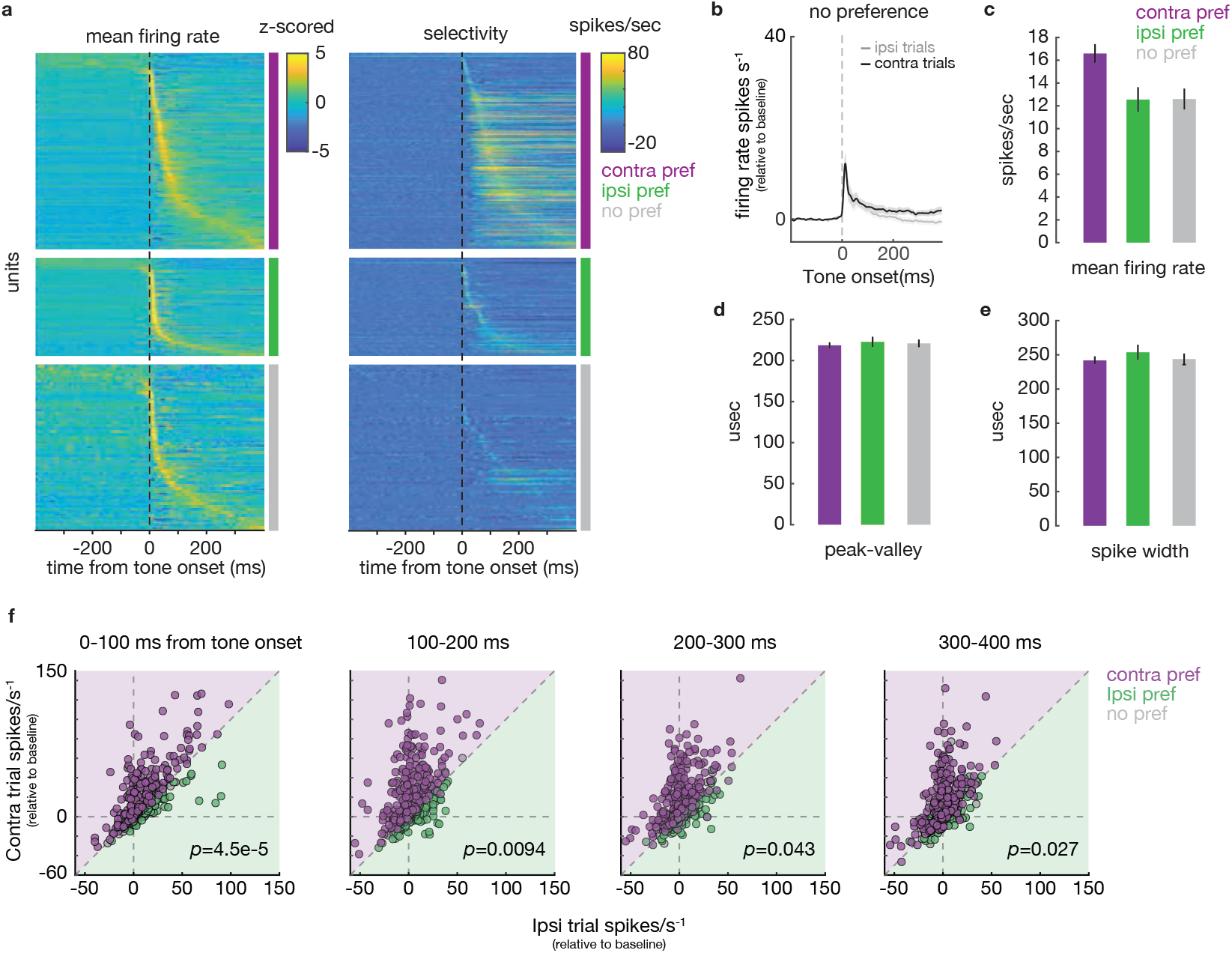
lSC activity and modulation by trial type. **a.** Mean firing rate (z-scored relative to firing during the ITI, left) and selectivity (spikes/s, right) of all lSC units. Each row shows data for a single unit, sorted by coding preference (right column for each panel). For each coding preference, units are sorted by the timing of peak firing relative to baseline. **b.** Firing rate (change from baseline in spikes/s) averaged across all non-selective (no preference) units during the first 100ms window after tone onset during contraversive (black) or ipsiversive (grey) correct trials. **c.** Mean firing rate (relative to baseline) calculated across sessions for each coding preference group. **d-e.** Peak-valley timing (d) and spike width (e) for waveforms of units in each coding group. No significant differences were observed between groups. **f.** Mean firing rates of individual units for contraversive vs. ipsiversive trials for different time bins relative to tone onset (as in Fig. 3h). P-values are shown (two-tailed *t*-test examining that activities in contraversive vs. ipsiversive trials differ).

**Extended Data Fig. 6.**
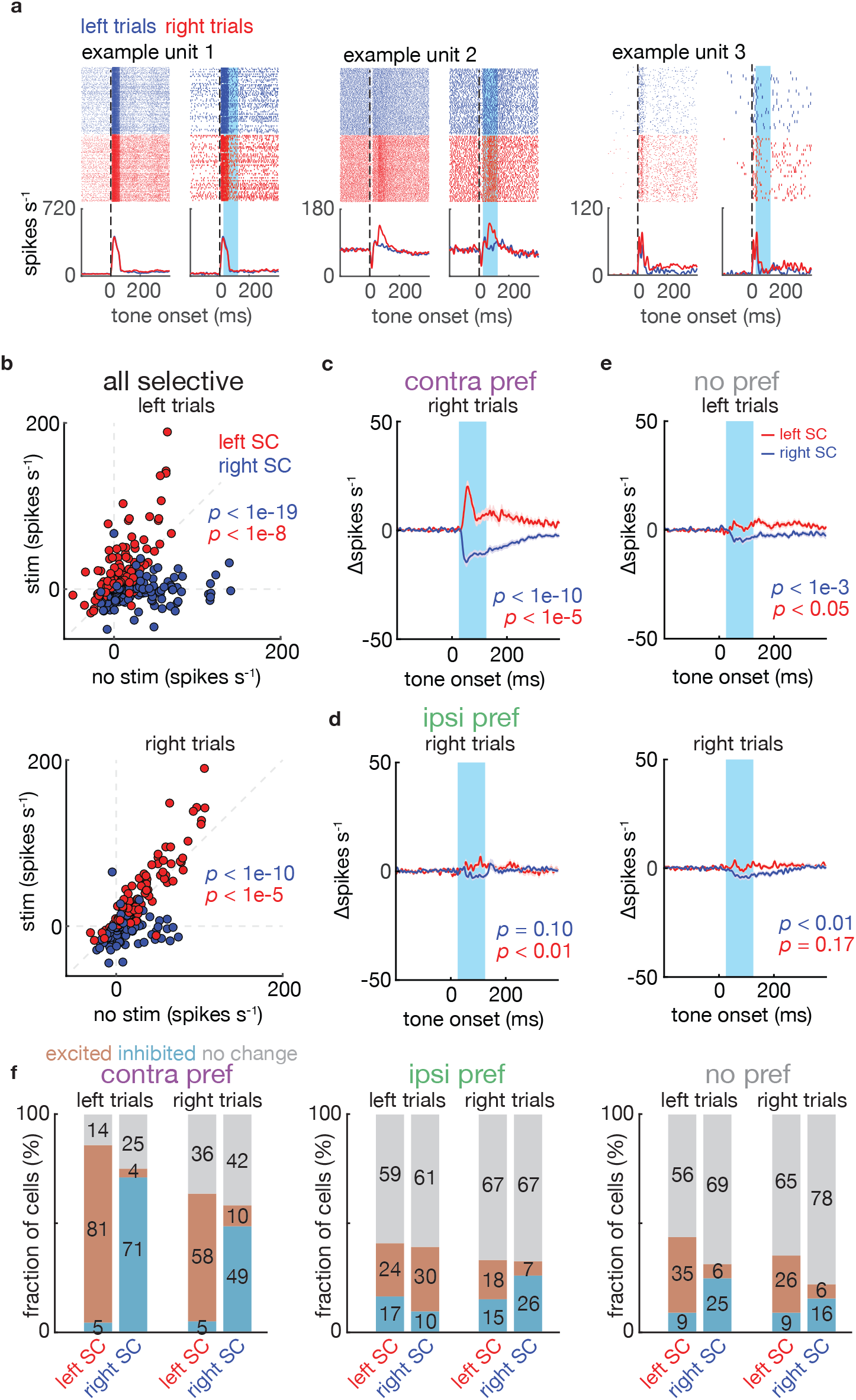
Detailed analysis of iSPNs modulation of lSC activity. **a.** Example units that were not significantly modulated by stimulation. Peri-stimulus histogram showing no stim trials (left) and stim trials (right, light blue=laser on) during left- (blue) and right- (red) cued trials. **b.** Average firing rates for each selective unit during the 100 ms laser-on period in stimulation (y-axis) and in the corresponding time in no stimulation (x-axis) trials (Main text Fig. 4d). Each dot represents a single unit recorded in either the left (red) or right (blue) SC. The optogenetic stimulation is always delivered in the right striatum. Firing rates are shown relative to baseline. P-value are shown for tests of significant modulation (two-tailed *t*-test). **c.** *left*, Changes in firing rates after stimulation during right-cued trials for contraversive preferring units (same analysis as Fig. 4d). **d.** As in panel **c** for ipsiversive preferring units. P-value are shown for tests of significant modulation (two-tailed *t*-test). **e.** As in panel **c** for units without significance lick direction tuning during left- (top) and right- (bottom) cued trials. **f.** Fractions of neurons that were excited, inhibited, or unchanged by optogenetic stimulation in left- and right-cued trials for contraversive-lick-preferring (*left*), ipsiversive-lick-preferring *(middle),* and untuned (no pref, *right)* groups (similar analysis as Fig. 4e) recorded in the left or right SC.

**Extended Data Fig. 7.**
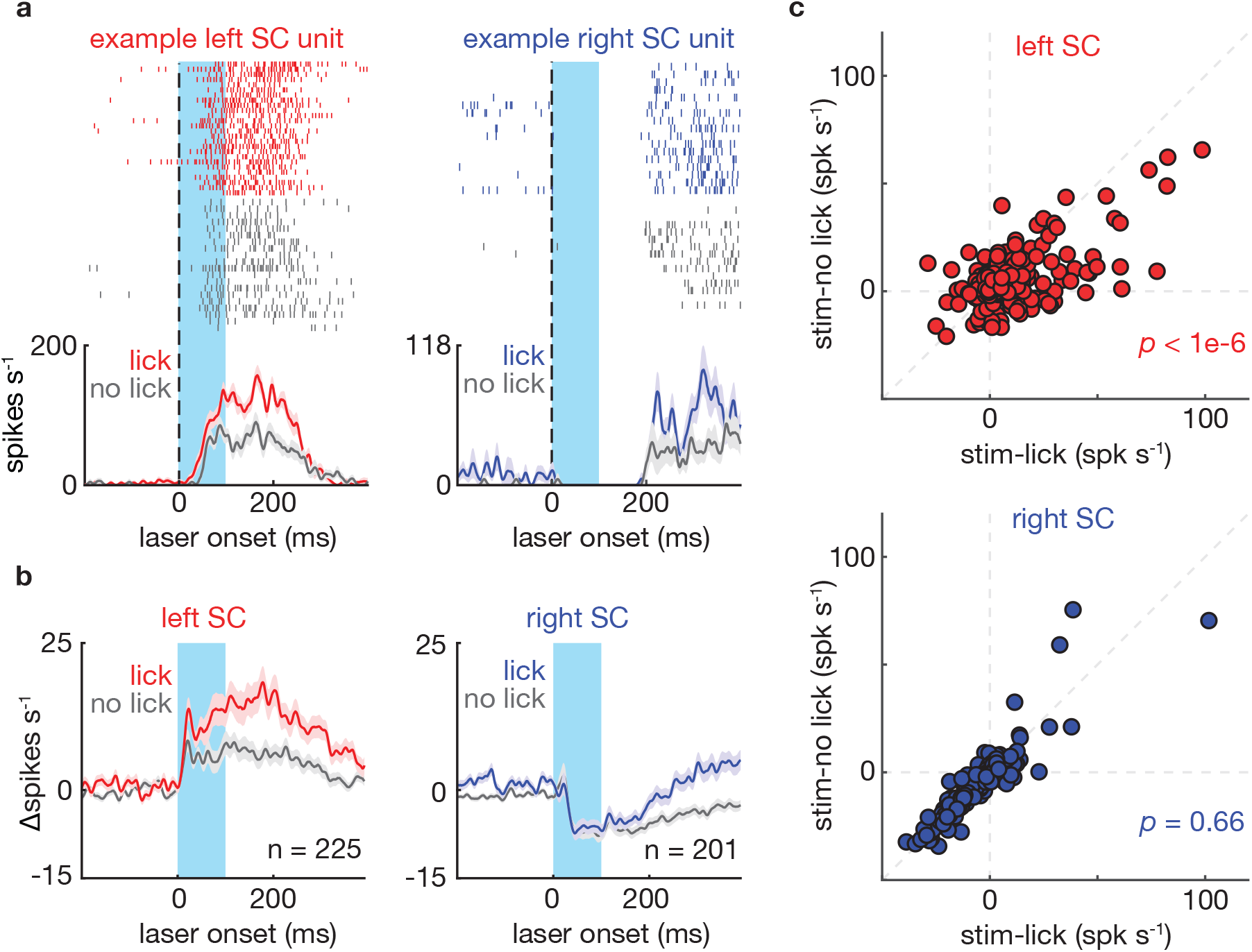
Effects of iSPNs stimulation during the ITI on SC activity. **a.** Example units recorded in the left SC (left panel) and right SC (right panel). Peri-stimulus histogram shows trials during which the stimulation induced licking (red/blue) or did not induce licking (grey). Firing rates shown as mean ± s.e.m across trials. **b.** Average changes in firing rate after stimulation (Δspikes s^-1^) in left SC (left panel) and right SC (right panel) grouped by behavioral outcome (red/blue=lick; grey=no lick). Firing rates shown as mean ± s.e.m across units (left SC: n=225; right SC: n=201). **c.** Average firing rates during the 100 ms stimulation window for stim-no lick trials (y-axis) vs. stim-lick trials (x-axis). Each dot represents a single unit. P-values show significance of modulation (two-tailed *t*-test).

**Extended Data Fig. 8.**
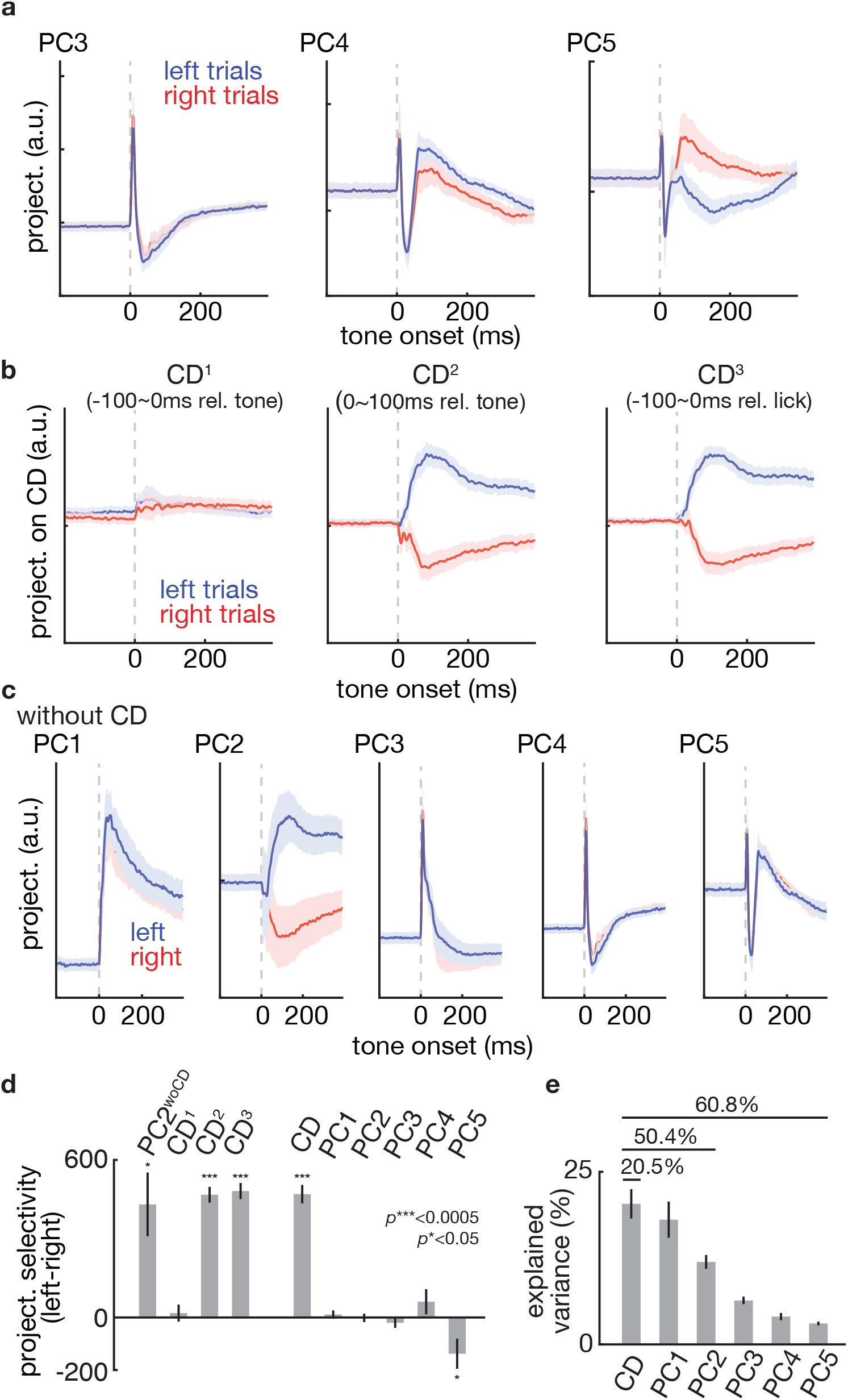
Detailed analysis of low dimensional projection of lSC activity. **a.** Activity projections onto PC3, PC4 and PC5 (see Main text, Fig. 5a-d). Left (blue) and right (red) trials are shown relative to tone onset. **b.** Comparison of different choices of coding direction vector. Coding direction was defined during −100~0ms window relative to tone onset (left, CD^1^), 0~100ms relative to tone (center, CD^2^) and 100~0ms relative to first lick (right, CD^3^) (see Methods). CD^1^ was used as a control. **c.** PCA on the original data (without first calculating and removing CD^2^ information as in Figure 5). Left/right lick (i.e. choice) information is found in PC2 (2^nd^ column). **d.** Left-right choice selectivity measured from the projection of the neural activity along the indicated axes. Selectivity measures how separable the trajectories are along the selected axis. The given P-values are for comparison by one sample two tailed *t*-test (P***<0.0005, P*<0.05). The trajectories are well-separable along different choice axes. PCs did not reliably discriminate trial type compared to CD (except for PC5). **e.** Explained variance along each dimension (see Methods). CD explained the most variance in the data (20.5 ± 2.3%). Explained variances for CD, CD+PC1+PC2, and CD+PC1~PC5 are shown. All error bars show bootstrapped standard error across units.

**Extended Data Fig. 9.**
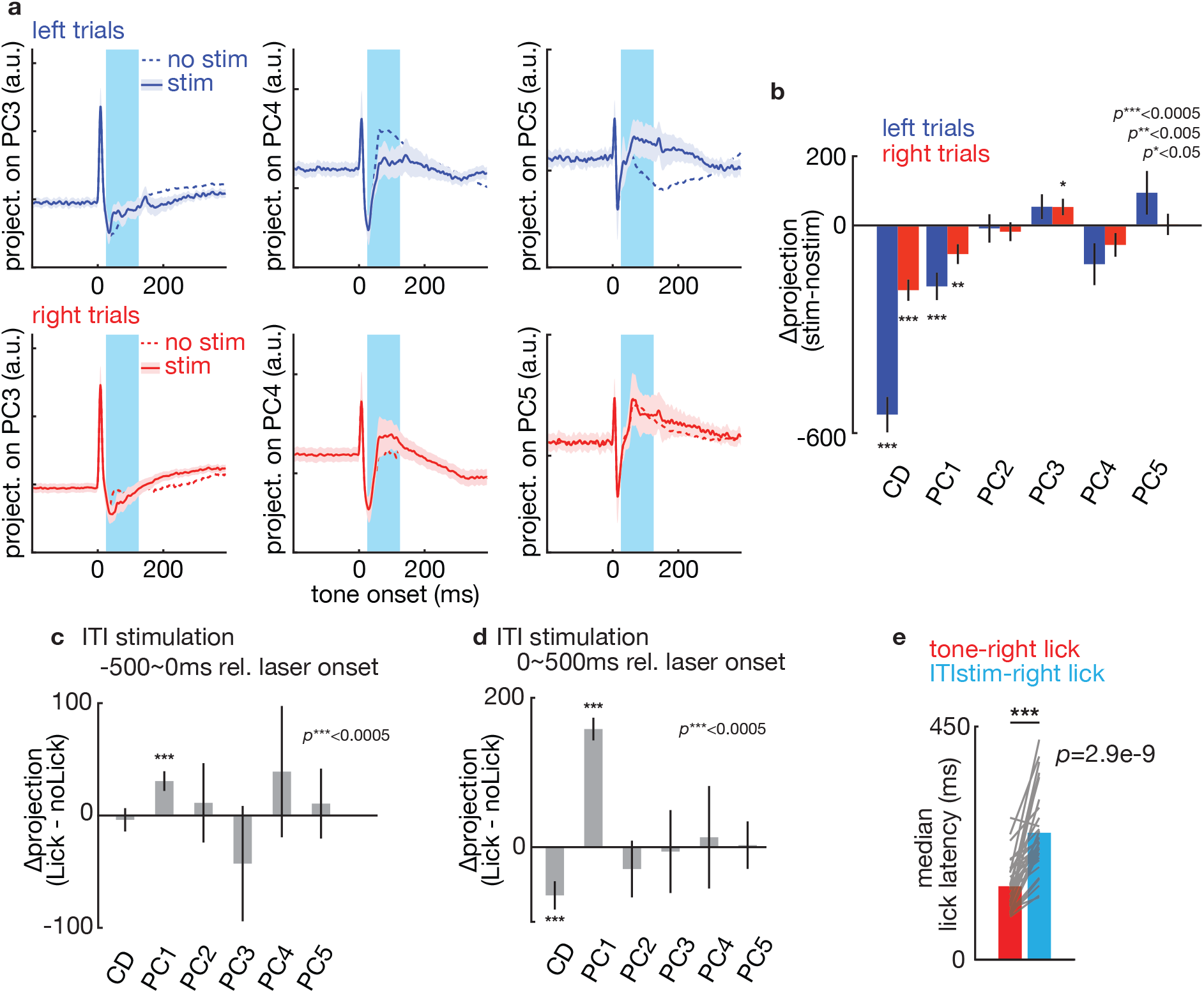
Detailed analysis of the effects of stimulation on the low-domensioal activity subspace. **a.** Projections of neural activity as a function of time relative to the tone onset shown along PC3 (left), PC4 (middle) and PC5 (right). Data are shown for left- (top, blue) and right- (bottom, red) cued trials. The dotted lines show activity in no stim trials and thick lines that in stim trials. **b.** Changes in activity during the stimulation window (100 ms) for each projection after stimulation (Δproject. modulation) along different dimensions during left- (blue) and right- (red) cued trials. Stimulation modulates activity the mostly along CD. P-values show significance of modulation (two-tailed t-test). **c-d.** Difference of activities along different directions in lick and no licks trials after ITI stimulation (see Main text, Fig. 5g). Panel **c** is for activity in the −500~0ms window relative to laser onset (i.e. 500 ms before the laser turned on) was used. PC1 discriminated whether the mice would lick after stimulation even before laser onset (P***<0.0005, two-tailed *t*-test). Panel **d** shows the same analysis for activity 0~500ms relative to laser onset (i.e. in the 500 ms after the laser turned on). PC1 best discriminated lick vs no-lick trials (P***<00005, two-tailed *t*-test). **e.** Median lick latency for right port licks triggered by the right-cue tone (control trials, red) or the optogenetic stimulation-induced during the ITI (blue). Each line shows averages for a single session (n=33). Only sessions with more than 5 stimulation induced licks were included. Licks induced by optogenetic stimulation during the ITI were slower than tone-triggered licks (P***<1e-8, two-tailed *t*-test). All error bars show bootstrapped standard error across units.

